# scPrediXcan integrates advances in deep learning and single-cell data into a powerful cell-type–specific transcriptome-wide association study framework

**DOI:** 10.1101/2024.11.11.623049

**Authors:** Yichao Zhou, Temidayo Adeluwa, Lisha Zhu, Sofia Salazar-Magaña, Sarah Sumner, Hyunki Kim, Saideep Gona, Festus Nyasimi, Rohit Kulkarni, Joseph Powell, Ravi Madduri, Boxiang Liu, Mengjie Chen, Hae Kyung Im

## Abstract

Transcriptome-wide association studies (TWAS) help identify disease causing genes, but often fail to pinpoint disease mechanisms at the cellular level because of the limited sample sizes and sparsity of cell-type–specific expression data. Here we propose scPrediXcan which integrates state-of-the-art deep learning approaches that predict epigenetic features from DNA sequences with the canonical TWAS framework. Our prediction approach, ctPred, predicts cell-type–specific expression with high accuracy and captures complex gene regulatory grammar that linear models overlook. Applied to type 2 diabetes and systemic lupus erythematosus, scPrediXcan outperformed the canonical TWAS framework by identifying more candidate causal genes, explaining more genome-wide association studies (GWAS) loci, and providing insights into the cellular specificity of TWAS hits. Overall, our results demonstrate that scPrediXcan represents a significant advance, promising to deepen our understanding of the cellular mechanisms underlying complex diseases.

## Introduction

Transcriptome-wide association studies (TWAS) are a class of methods that nominate candidate causal genes for complex traits and diseases by determining associations between predicted gene expression and phenotype^1–3^. Canonical TWAS approaches train a gene expression prediction model using tissue-level gene expression from a reference panel of at least 100 individuals. While TWAS has been successfully applied to various tissues and traits, providing candidate causal gene lists^4,5^, it is limited by the mismatch between available expression panels and disease-relevant cell types or states. Tissues with extensive expression panels (e.g., whole blood or lymphoblastoid cell lines) are commonly employed to maximize power, but recent studies suggest that context-specific regulation of gene expression is more relevant for disease^3,6,7^. This is especially true when rare cell types that are underrepresented in conventional bulk-tissue expression data drive disease onset. Therefore, a TWAS framework with a model that can predict expression from disease-relevant tissues and/or cell types would be of great value for uncovering genes involved in the trait etiology. Although our understanding of cellular heterogeneity and cell-type-specific regulatory patterns has improved dramatically with recent advances in single-cell RNA sequencing (scRNA-seq)^8^, single-cell data is simply not available on the scale required by canonical TWAS frameworks to ensure optimal accuracy and power across contexts.

Recent advancements in deep learning models that predict molecular features based on sequence data present a promising solution. While existing deep learning models for gene expression predictions are trained on bulk RNA-seq data and are constrained to tissue-level predictions, models predicting molecular features (e.g., epigenome) are trained using only a reference genome, thus eliminating the need for population-level data^9,10,11^. To overcome this, we employed transfer learning to develop a cell-type–specific gene expression prediction model, ctPred, trained from pseudo-bulk scRNA-seq data. This model leverages knowledge from pre-trained sequence-to-molecular features models, enhancing convergence, reducing overfitting, and improving overall performance. Specifically, we leveraged this advantage by utilizing the state-of-the-art sequence-to-epigenomics model, Enformer, as a feature extractor. We employed Enformer-predicted features via transfer learning to develop a cell-type–specific gene expression prediction model, **ctPred**, trained from pseudo-bulk scRNA-seq data. By building on knowledge from pre-trained sequence-to-epigenomics models, ctPred enhances convergence, reduces overfitting, and improves overall performance. We further developed **single-cell PrediXcan (scPrediXcan),** a cell-type-level TWAS framework, by leveraging ctPred to perform TWAS using single-cell data. While ctPred is theoretically capable of predicting context-specific, in-silico expression data at the individual level for TWAS using GWAS data, the computational demands remain prohibitive given the current scale of GWAS (i.e., hundreds of thousands of individuals). Additionally, access to individual-level GWAS data is often difficult to obtain. To overcome these challenges, we introduced a SNP-based linear version of ctPred, termed **ℓ-ctPred**, which is derived from genotype data alongside a ctPred-predicted, in-silico expression reference panel. Using ℓ-ctPred’s weights, we can conduct association tests between genes and diseases using only summary statistics from GWAS data, thereby enabling a TWAS that is fundamentally based on single-cell data.

We evaluated scPrediXcan by applying it to two diseases, type 2 diabetes (T2D) and systemic lupus erythematosus (SLE). Our comparison of scPrediXcan’s performance against canonical TWAS models trained using the same datasets reveals that scPrediXcan significantly outperforms the canonical frameworks. It identifies a larger number of candidate causal genes, explains more GWAS loci, and provides more detailed insights into the cellular specificity of the TWAS findings. These results highlight scPrediXcan’s substantial improvement in both the accuracy and relevance of gene nominations within TWAS, demonstrating its promising potential to advance our understanding of the cellular mechanisms that underpin complex diseases.

## Results

### Overview of scPrediXcan framework

The scPrediXcan framework consists of three key steps (Fig. 1, supplementary fig. 2). First, we trained a model to predict gene expression from epigenomic data. Second, we linearized this deep-learning model into a SNP-based elastic net model, which can be used for association tests using GWAS summary statistics. Finally, we tested associations between genes and the trait of interest.

**Figure 1.**
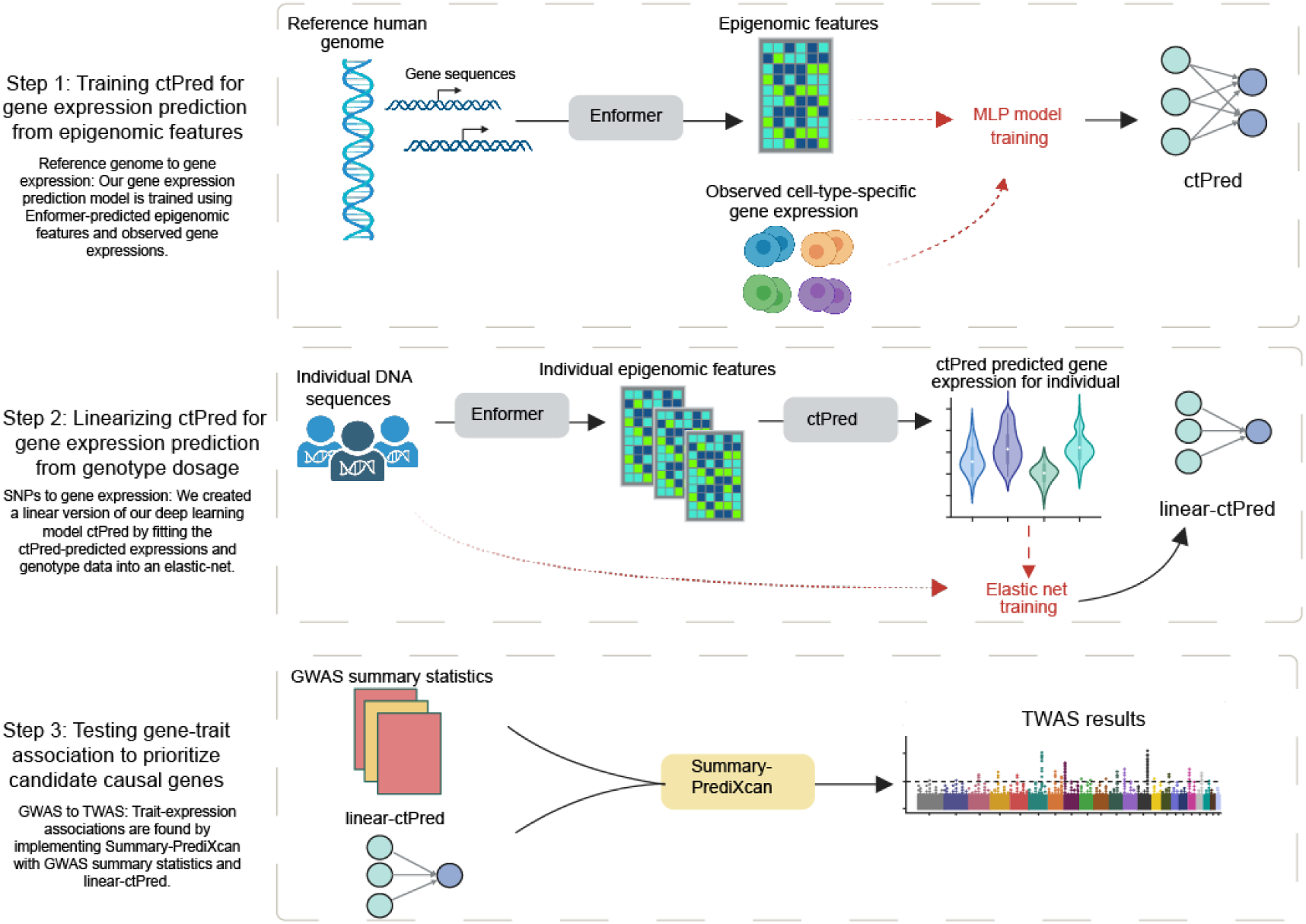
Overview of scPrediXcan framework.

In the first step, we established a method to predict the cell-type–specific gene expression from the DNA sequences. Because it is challenging for the model to learn genomic grammar by training directly on the highly sparse single-cell data, we utilized the genomic regulation insights gained from the state-of-the-art sequence-to-epigenomics model trained on bulk-level data, Enformer^9^. We used Enformer as a feature extractor and, through transfer learning, we trained ctPred—a lightweight, four-layer multi-layer perceptron (MLP) designed for predicting cell-type–specific gene expression at the single-cell pseudobulk level.

Despite ctPred’s ability to predict individual cell-type–specific gene expression from gene sequences, the computational expense of such predictions for TWAS association tests across large GWAS cohorts remains high, and access to individual-level data is limited. We addressed this in our workflow’s second step by transforming the deep learning model into a linear form, creating a SNP-based elastic net version of ctPred, termed ℓ-ctPred. This version can predict gene expression for specific cell types from SNP dosages. The weights for ℓ-ctPred are stored in a database, eliminating the need for end-users to repeat the training steps.

The final step of our workflow tests for associations between genes and traits or diseases at the cell-type level using S-PrediXcan^12^. This step employs the weights from ℓ-ctPred along with GWAS summary statistics to estimate gene effect sizes on the trait and to compute p-values, thus prioritizing putative causal genes.

### ctPred accurately predicts single-cell pseudobulk gene expression across the genome in diverse cell types and datasets

To develop prediction models for different cell contexts and evaluate their performance, we trained ctPred and tested its prediction performance on three scRNAseq datasets separately: OneK1K dataset^13^, a T2D islet dataset, and a subset of the Tabula Sapiens dataset^14^. The OneK1K dataset includes 29 cell types from 982 individuals, the T2D islet dataset has 11 cell types from 29 individuals, and the Tabula Sapiens dataset contains more than 400 cell types from 14 different organs of 15 individuals. To conserve computational resources, we analyzed a subset of the Tabula Sapiens dataset comprising 6 cell types from 15 individuals.

For each dataset, we tailored ctPred models to individual cell types. To avoid data leakage due to sequence overlaps between genes, we randomly divided the datasets by chromosomes into training, validation, and test sets (Fig. 2a). We then evaluated the models’ performance by calculating Pearson correlations between predicted and observed pseudobulk gene expressions across the test sets (Fig. 2b). For example, in the OneK1K dataset, Pearson correlations for all cell types exceed 0.8, with the highest at 0.892 in CD4+ alpha-beta T cells and the lowest at 0.836 in erythrocytes (Fig. 2c). indicating that ctPred accurately predicts gene expression across diverse cell types.

**Figure 2:**
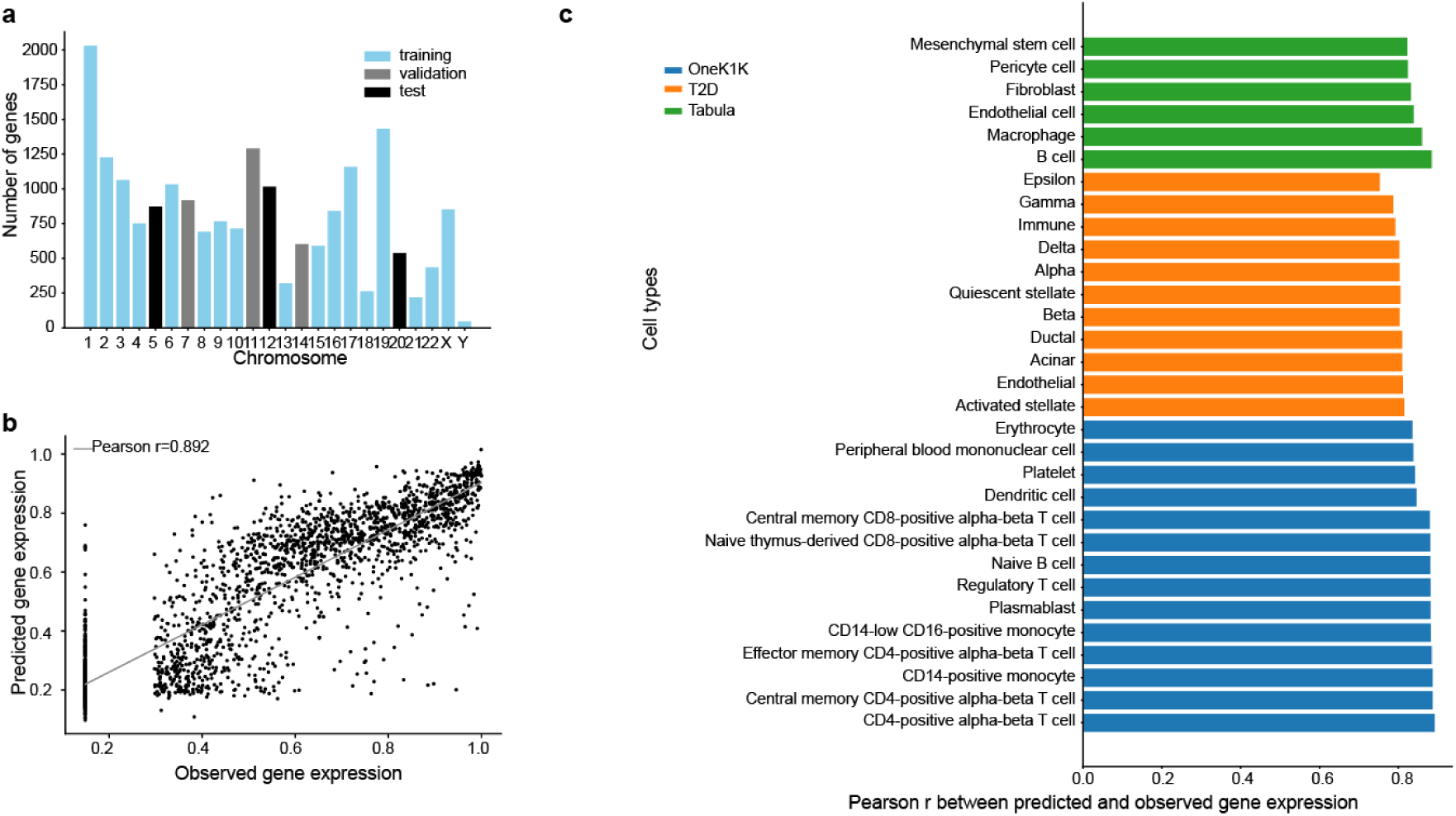
ctPred predicts cell type-specific gene expression across the genome. **a**) Data splitting by chromosomes for ctPred training. **b**) Scatter plot of the predicted gene expression and observed gene expression for each gene in a CD4+ alpha-beta T cell from OneK1K dataset. **c**) Bar plot of Pearson correlations between predicted gene expression and observed gene expression in various cell types from OneK1K, T2D, and Tabula Sapiens datasets. Results for all cell types are listed in supplementary table 3.

Further testing the model’s robustness, we validated ctPred on 11 cell types from the T2D dataset and 6 from the Tabula Sapiens subset. In the T2D dataset, Pearson correlations range from 0.753 in epsilon cells to 0.815 in activated stellate cells. In the selected Tabula Sapiens cell types, correlations range from 0.823 in mesenchymal stem cells to 0.885 in B cells, showcasing the model’s effectiveness across various datasets and cellular contexts.

During our project, we learned of the parallel development of a module within seq2cells^15^ called emb2cell, which specifically focuses on single-cell pseudobulk gene expression prediction. Similar to our ctPred, emb2cell uses embeddings for prediction, but while ctPred utilizes the 5313-dimension output from Enformer, emb2cell leverages 3072-dimension intermediate embeddings to train a two-layer MLP model. A key distinction between the models is that ctPred is much more parameter-efficient, containing approximately 0.4M parameters compared to emb2cell’s over 60M. We evaluated both models using CD4+ T cell scRNA-seq data, adhering to the training, validation, and testing partitions specified in the seq2cells publication. ctPred achieves a Pearson correlation of 0.787 across genes (supplementary fig. 1), significantly outperforming the 0.666 correlation reported for emb2cell.

The minimum number of cells per cell type was 125 in OneK1K, 212 in T2D, and 4,539 in Tabula Sapiens. In general, prediction performance improved with an increasing number of available cells within each cell type, underscoring the benefits of our approach, where reads are aggregated across individuals (Supplementary Figures 7a, e, and i). This advantage becomes even more evident in the following section, where we assess performance across individuals.

Overall, across 46 cell types and three datasets, ctPred not only achieves Pearson correlations ranging from 0.753 to 0.892, but also surpasses emb2cell on the CD4+ T cell dataset, despite having 150 times fewer parameters. This underscores ctPred’s efficiency, accuracy, and robustness in predicting cell-type–specific gene expression.

### ctPred outperforms SNP-based predictors used in canonical TWAS for predicting cell-type–level gene expression across individuals

To assess ctPred’s performance across individuals—crucial for TWAS aiming to discern gene expression changes between patients and healthy controls—we compared it with a SNP-based approach used in canonical TWAS. This SNP-based method, which we refer to as pseudobulk elastic net (PEN), is trained on the same observed single-cell data as ctPred and should not to be confused with ℓ-ctPred, which also uses elastic net but is based on in-silico gene expression predictions.

We found that PEN fails to converge for the majority of genes attempted, yielding predictors for only ~3.5% of expressed genes, whereas the ctPred approach yields predictors for all expressed genes (~19,000). Specifically, PEN yields 318–434 (median = 354) and 504–1784 (median = 707) across cell types in T2D and OneK1K, respectively. Fig. 3a provides examples of ctPred-predicted versus observed gene expressions, highlighting some genes missed by the PEN model.

**Figure 3.**
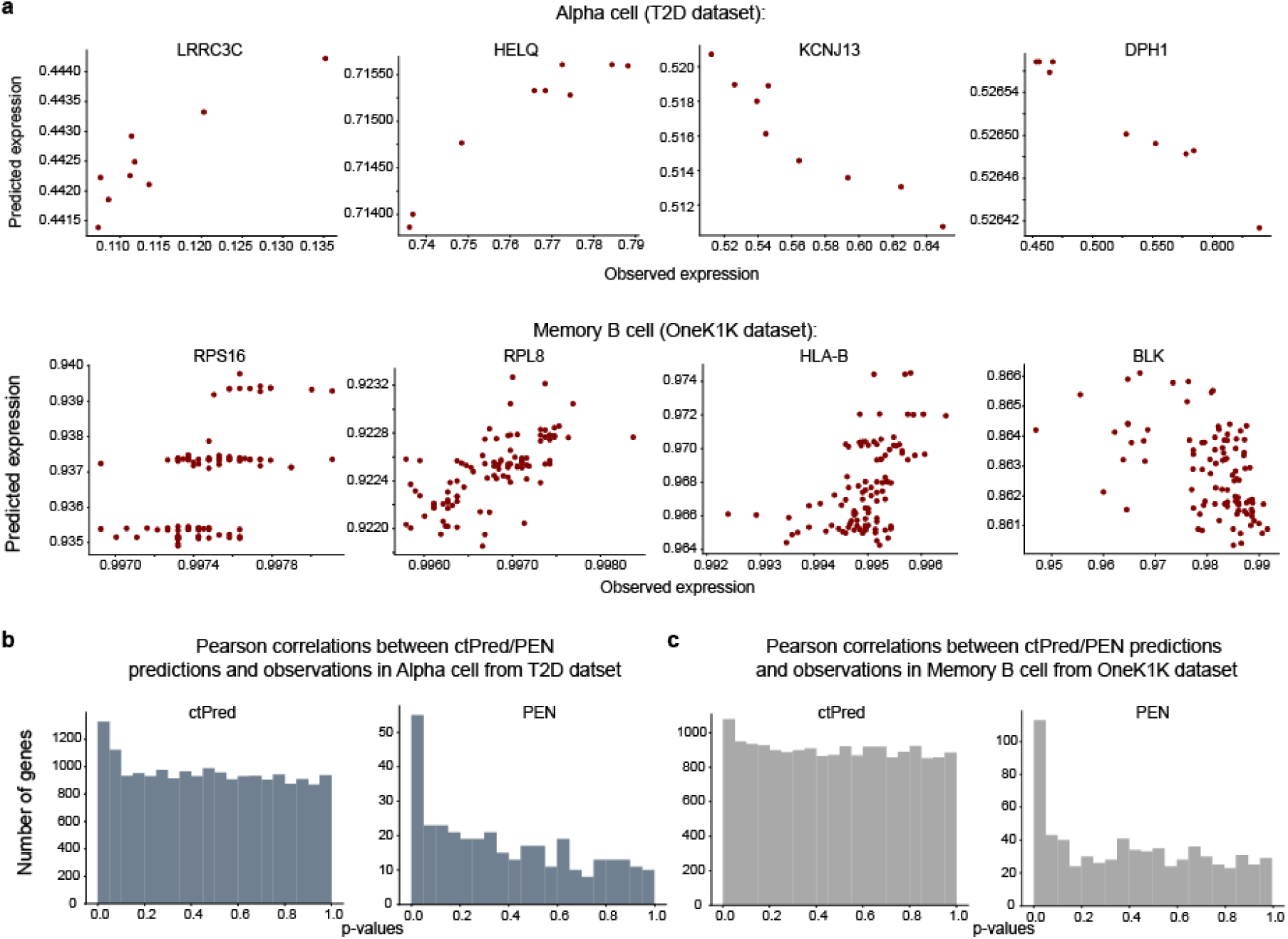
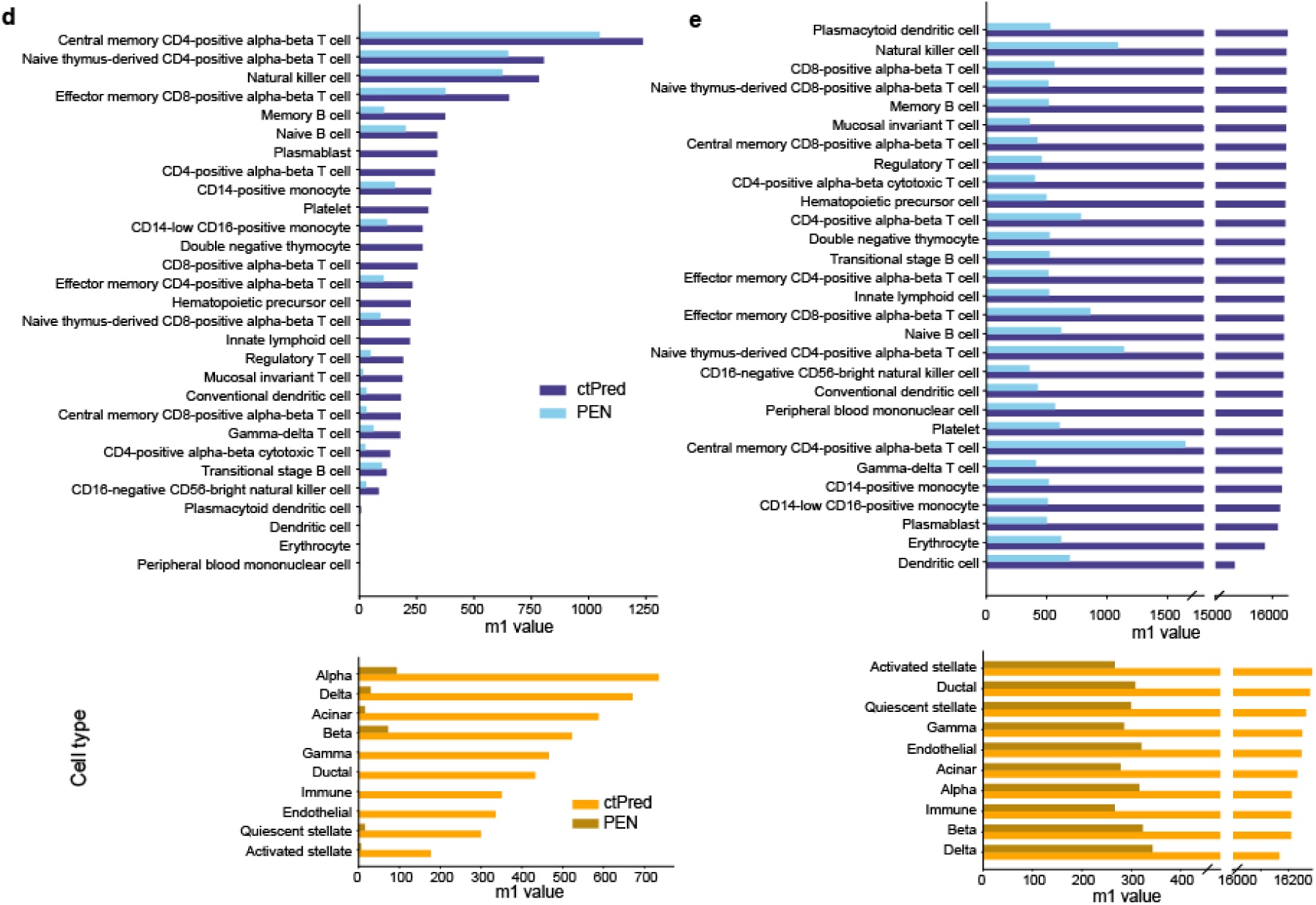
ctPred predicts cell type-specific gene expressions across individuals. **a**) Scatter plot of ctPred predicted-gene expression and observed gene expression for representative genes missed by PEN in Alpha cell and Memory B cell in the test set. **b**) Histogram of the distribution of Pearson correlation p-values between model-predicted gene expression and observed gene expression in Alpha cell from T2D dataset. **c**) Histogram of the distribution of Pearson correlation p-values between model-predicted gene expression and observed gene expression in Memory B cell from OneK1K dataset. **d**) Bar plot of m1 values between predicted gene expression and observed gene expression of ctPred/PEN in different cell types. **e**) Bar plot of m1 values between ctPred/PEN-predicted gene expression and bulk GReX of in different cell types.

Next, we assessed the correlation between the predicted and observed gene expression across individuals to evaluate predictor performance. Due to the known correlation sign inconsistency in Enformer^16,17^, we utilized the p-value of correlation as the primary performance metric (see Discussion). The histograms of p-values for correlations in alpha cells and memory B cells are shown in Fig. 3b-c. However, given the substantial difference in the number of genes predicted by each model, caution must be taken to ensure a fair comparison. Since convergence and prediction performance are intertwined, focusing solely on genes that converge in PEN would introduce bias. One robust metric is the number of ‘true positive genes’ (i.e, m1= π* #genes), estimated using the q-value framework^18^ and represents genes with p-values deviating from a uniform distribution (supplementary fig. 4). Our ctPred approach consistently identifies a larger number of true positive genes compared to the traditional PEN method when validated against observed expression, though the total number of genes conclusively associated with observed expression is modest, ranging from 0 to 1236, with a median of 275 (Fig. 3d).

Since our primary objective is to elucidate how GWAS variants influence phenotypes through the regulation of molecular phenotypes like cell-type–specific gene expression (i.e., genetically regulated expression, GReX), we also evaluated the performance of ctPred and PEN using GTEx-trained bulk predictors^19^ as proxies for the genetically regulated component of expression (GTEx GReX). We predicted gene expression in 400 European individuals using 1000G cohort^20^ genotype data and GTEx-trained weights, which we compared against both ctPred and PEN predictions in the same cohort. We calculated correlations, derived p-values, and estimated π_1_ to assess the proportion of genes for each model genuinely associated with GTEx GReX proxies. This analysis primarily identifies associations between components of GReX shared across cell types and bulk tissues, suggesting that the ability to predict shared regulation may indicate potential in predicting cell-type–specific components, despite data limitations for direct testing. We evaluated the correlation of our cell-type expression predictors with GTEx GReX across 49 tissues, aggregating results into a single p-value per gene per cell type using the ACAT method^21^. Approximately 19,000 genes were tested using ctPred predictors, while PEN predictors were used to test between 318 and 1784 genes, constrained by the limited predictors this approach yields. Using the π_1_ statistic, we estimated 95.3%–96.1% (median = 95.7%) of ctPred-predicted genes and 59.2%–95.5% (median = 77.8%) of PEN-predicted genes are truly correlated with GTEx GReX. The disparity between model performance is even more stark when considering the number of truly correlated genes: ctPred predicts 15,339–16,277 (median = 16,206) truly correlated genes, while PEN only predicts 277–1,646 (median = 458).

These findings confirm that ctPred more effectively predicts shared regulation than PEN and suggests that ctPred may also better predict cell-type–specific regulation. Given ctPred’s robust performance, we decided to include all genes predicted by ctPred in our phenotype association tests. According to recent analysis^22^, incorporating genes not associated with GReX does not compromise the type I error rate, provided that the appropriate adjustments for multiple testing are made.

Furthermore, our comparison of performance across individuals and the number of cells per cell type reveals a similar trend to that observed for performance across genes. As shown in Supplementary Fig. 7, the number of true positive genes per cell type increases with the total number of available cells. The canonical TWAS, which relies on variation across individuals, is more sensitive to a low number of cells compared to ctPred, which gains robustness by aggregating cells across individuals.

### Linear-ctPred enables large-scale context-specific TWAS

Having developed reliable context-specific prediction models, we are theoretically equipped to conduct TWAS using single-cell informed expression levels. However, the computational burden using Enformer and ctPred at the scale required is significant; we estimate predicting gene expression for 500 individuals across 20,000 genes would require ~2,700 GPU-hours. Additionally, individualized data (e.g., raw GWAS data) is difficult to access for most diseases. An efficient alternative is to use readily accessible GWAS summary statistics for association studies. S-PrediXcan is the commonly used method for performing TWAS using GWAS summary statistics^12^; however, it requires linear gene expression predictors, meaning ctPred is incompatible with the framework. To address this incompatibility, we created an in silico reference dataset using ctPred predictions for several hundred individuals and fitted a SNP-based elastic net model to this data. This resulting model, ℓ-ctPred, takes individual genotype data and yields predicted expression levels using linear combinations of SNP dosages (Fig 1, step 2).

We linearized ctPred models for 40 cell types from T2D and OneK1K datasets using genotype data of 462 European individuals from the 1000G project and stored the weights for downstream association analysis. We used European ancestry data to better match the ancestry of available GWAS. We validated linearization efficiency by calculating the 10-fold cross-validated Spearman correlations between predictions from ctPred and ℓ-ctPred across all genes (supplementary fig. 5). For all cell types, the median correlation is above 0.83, indicating that ℓ-ctPred robustly approximates ctPred (Fig. 4). Thus, we integrated ℓ-ctPred as the gene expression predictor in the scPrediXcan framework.

**Figure 4:**
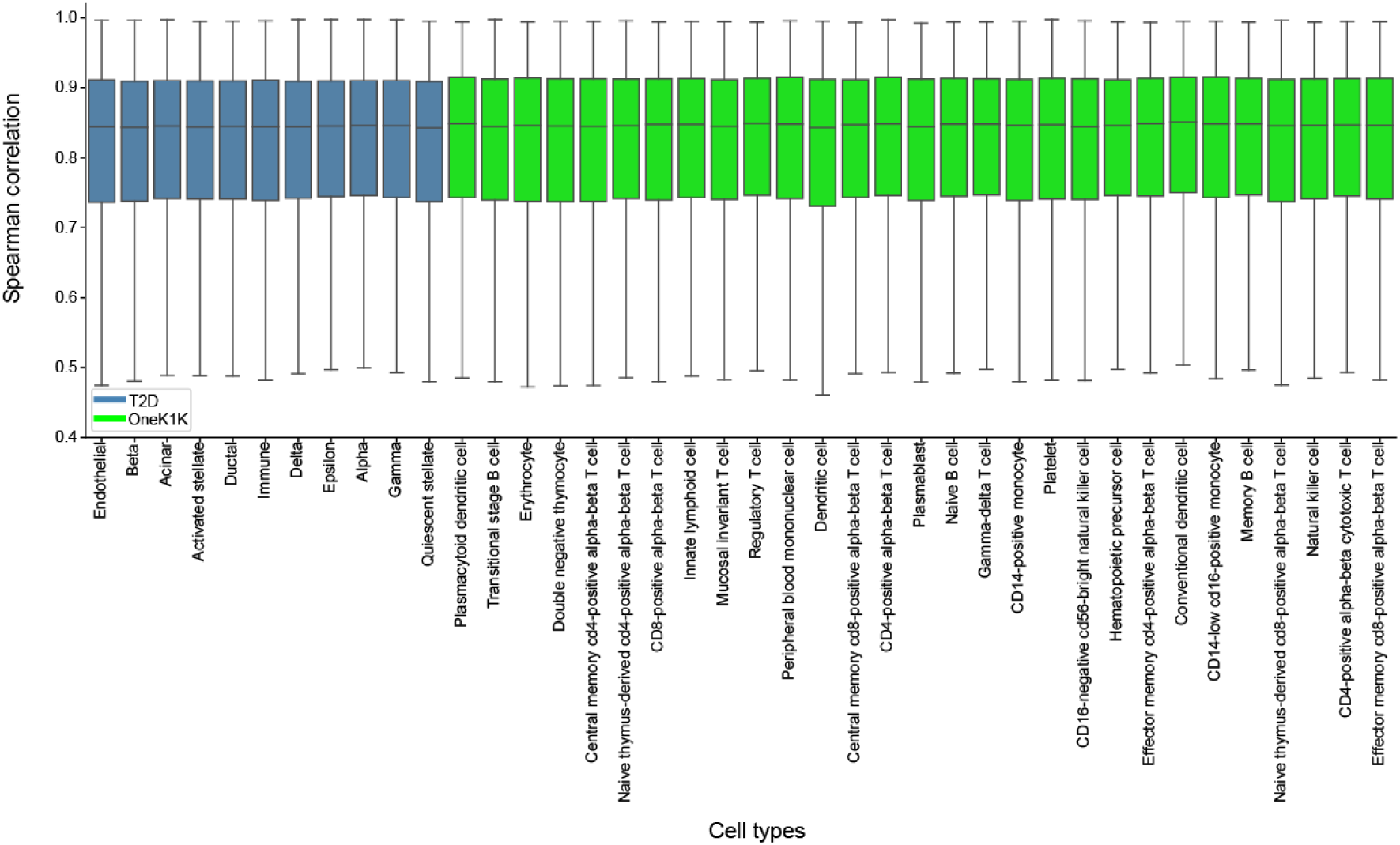
l-ctPred approximates crPred with high concordance. Bar plot of Spearman correlations between ctPred predictions and ℓ-ctPred predictions across individuals in 40 different cell types from T2D and OneK1K datasets.

While both the canonical TWAS gene expression predictors and ℓ-ctPred use a SNP-based elastic net model, ℓ-ctPred has a significantly higher number of genes converged during the model training. For example, for the memory B cell type from OneK1K dataset, the canonical SNP-based model trained on observed expression data achieved convergence for 340 genes with a training sample size of 800 individuals and required both genotype and scRNA-seq data. In contrast, ℓ-ctPred achieved convergence for 16,892 genes with a training sample size of only 462 individuals using solely genotype data. The stark difference in the number of converging genes arises because the canonical TWAS prediction model is based on observed data, whereas ℓ-ctPred relies on in-silico ctPred-predicted gene expression data. The in-silico data contains only the genetic component of expression, while the observed gene expression includes both genetic and environmental components, making ℓ-ctPred less vulnerable to the noise and sparsity present in observed scRNA-seq data. Furthermore, ℓ-ctPred is not limited by the small sample sizes associated with observed scRNA-seq data, as it can predict more individual expression as needed from the given genotype data.

### scPrediXcan enables TWAS at the single-cell pseudobulk level for type 2 diabetes and outperforms canonical TWAS methods

Our context-specific TWAS framework, scPrediXcan, utilizes cell-type–specific ℓ-ctPred models as the predictive component within the S-PrediXcan framework to explore gene–disease associations at the cell-type level (Fig. 1, step 3). To demonstrate the efficacy of scPrediXcan in identifying candidate causal genes, we conducted comparisons involving several benchmarks: 1) a cell-type–specific canonical TWAS method (TWAS-pseudobulk) that uses the same scRNA-seq pseudobulk data as scPrediXcan (i.e., PEN predictors); 2) a tissue-level canonical TWAS method (TWAS-bulk) that relies on bulk RNA-seq data; and 3) other gene prioritization methods for T2D (Methods).

scPrediXcan identifies a larger number of candidate T2D-associated genes across more GWAS loci compared to both TWAS-pseudobulk and TWAS-bulk. Specifically, using scPrediXcan, we identified 222 candidate causal genes across 108 different linkage disequilibrium (LD) blocks from a total of 1703 pre-defined approximately independent LD blocks^23^. In contrast, we identified only 12 candidate genes across 11 LD blocks with TWAS-pseudobulk, and 111 genes across 64 LD blocks with TWAS-bulk. Representative results for all three frameworks are shown for beta cells (scPrediXcan and TWAS-pseudobulk) and pancreas (TWAS-bulk) in Figure 5a. The full set of association statistics are in Supplementary tables 4–14.

**Figure 5:**
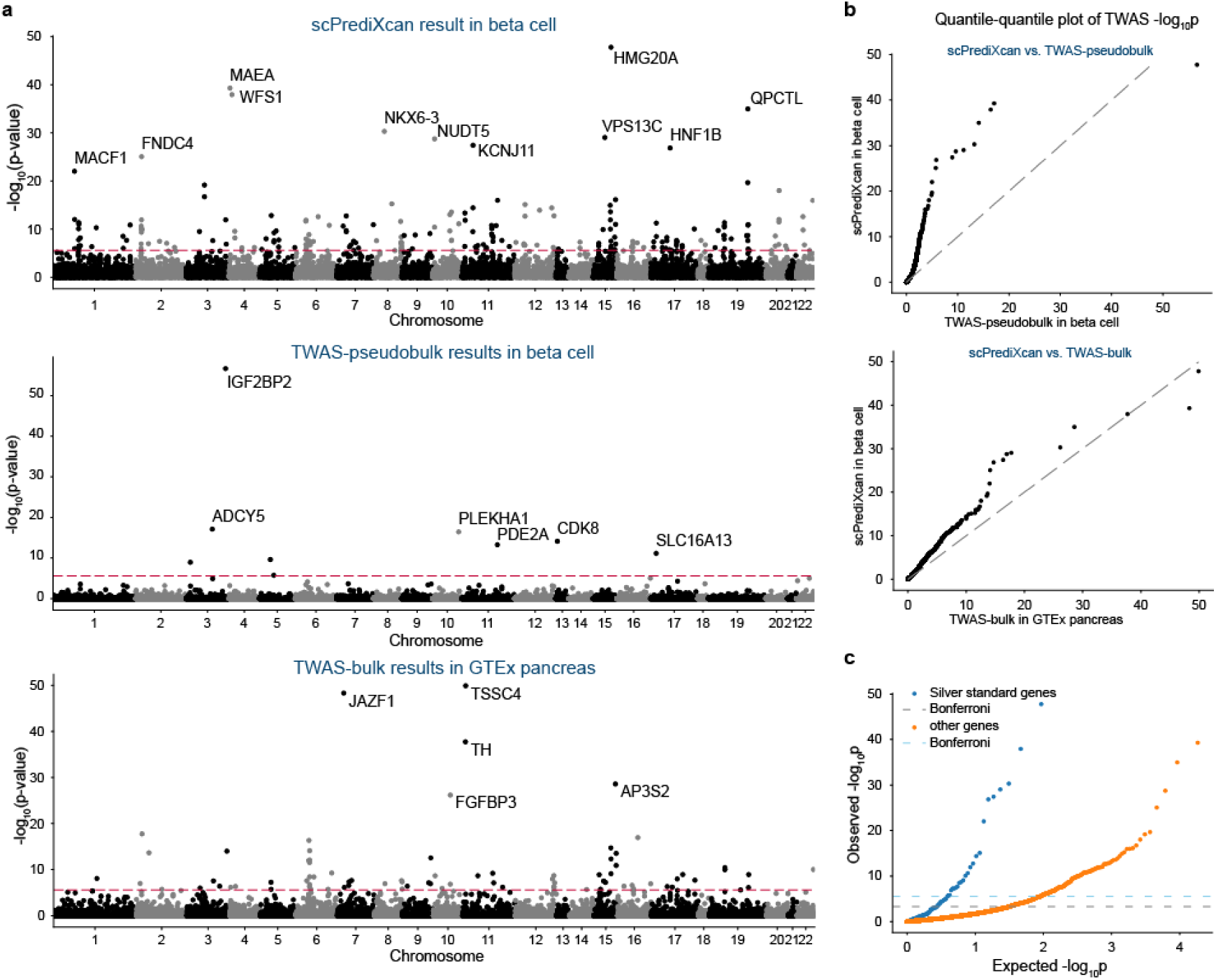
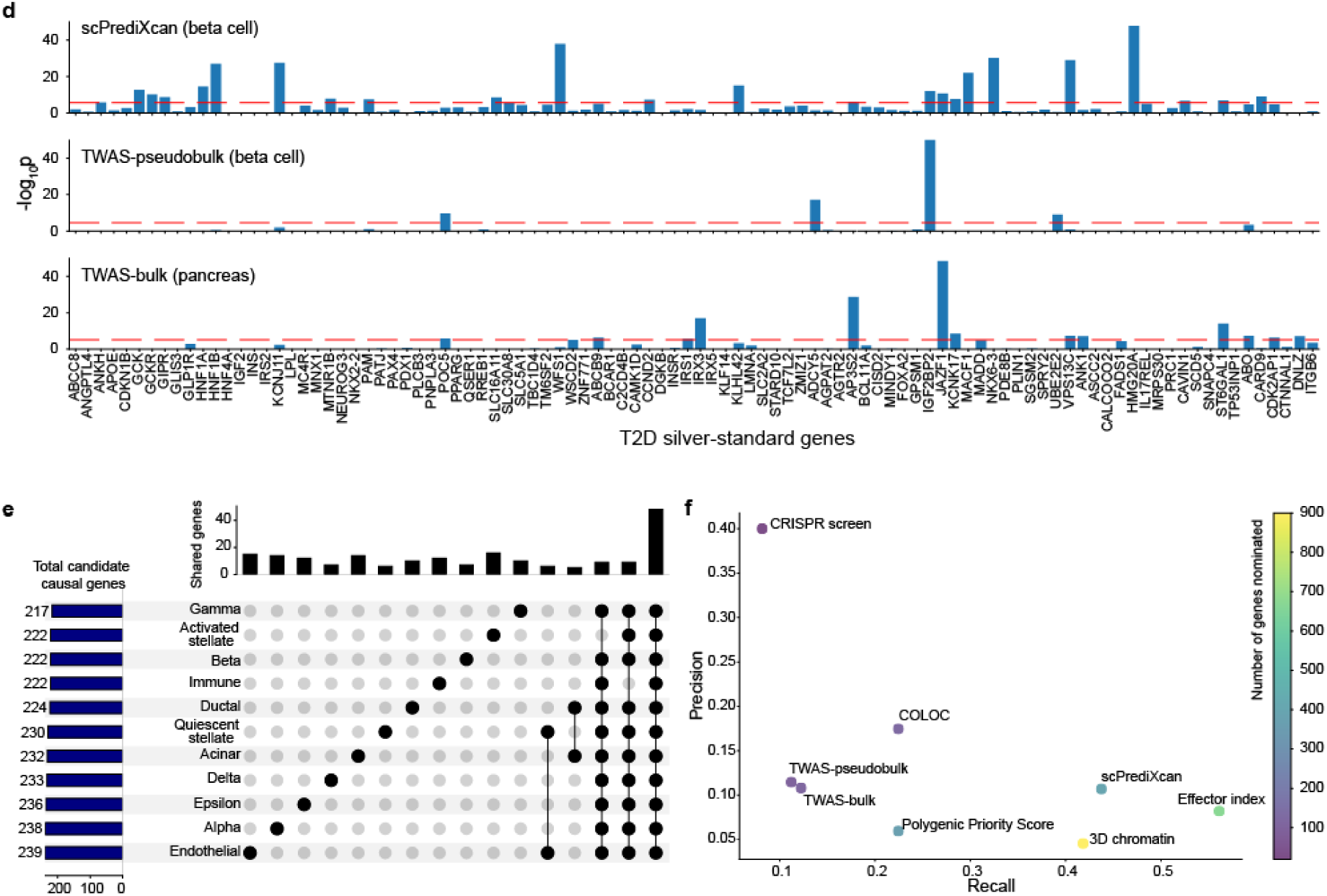
scPrediXcan in type 2 diabetes outperforms canonical TWAS methods. **a**) Manhattan plots of T2D TWAS results for different frameworks. Top: scPrediXcan in beta cell from T2D dataset. Middle: TWAS-pseudobulk in beta cell from T2D dataset, Bottom: TWAS-bulk in pancreas tissue from GTEx dataset. The red dashed lines are Bonferroni-corrected thresholds (p<0.05/number of genes in the association study). **b**) QQ-plot of TWAS p-values in T2D between frameworks. **c**) QQ-plot of TWAS p-values in T2D of scPrediXcan against the null distribution (i.e., uniform distribution). Blue: silver standard genes, Orange: other genes. Dashed lines: Bonferroni-corrected thresholds. **d**) Bar plot of TWAS −log10(p) of T2D silver standard genes in different frameworks. Top: scPrediXcan in beta cell from T2D dataset. Middle: TWAS-pseudobulk in beta cell from T2D dataset. Bottom: TWAS-bulk in pancreas tissue from GTEx dataset. **e**) UpSet plot of scPrediXcan-nominated candidate causal genes for T2D in different cell types. **f**) Scatter plot of precision and recall of different gene-prioritization methods for T2D causal gene nomination.

Further, we evaluated the statistical significance of these findings by comparing the TWAS p-values of scPrediXcan against those from TWAS-pseudobulk and TWAS-bulk through a quantile–quantile plot (QQ-plot, Fig. 5b). Considering that ℓ-ctPred achieves convergence for significantly more genes than the SNP-based models used in the other two frameworks, we used a uniform distribution of p-values to represent genes absent in the canonical approaches, ensuring a comprehensive comparison. The QQ-plot clearly demonstrates that scPrediXcan outperforms the canonical TWAS frameworks, consistently showing statistically lower p-values for identified TWAS hits.

To evaluate scPrediXcan’s efficacy in identifying causal genes for type 2 diabetes (T2D), we utilized a curated list of T2D-associated genes from the Common Metabolic Diseases Knowledge Portal (CMDKP) database^24^, which we call T2D ‘silver-standard’ genes (Supplementary table 45). We compared the scPrediXcan p-values for these genes against those of the remaining genes predicted by ℓ-ctPred. A QQ-plot against a uniform p-value distribution (Fig 5c) demonstrates that the silver-standard genes have significantly lower p-values, affirming that scPrediXcan accurately identifies genes truly associated with T2D. We also compared these results with those from canonical TWAS frameworks; we show representative results for beta cells (Fig 5d). Among 98 silver-standard genes, scPrediXcan has 24 Bonferroni-corrected significant genes (p < 2.7 × 10^−6^), whereas TWAS-pseudobulk and TWAS-bulk identify only 4 (p < 2.8 × 10^−6^) and 13 (p < 8.5 × 10^−6^) significant genes, respectively. Notably, both scPrediXcan and TWAS-pseudobulk recognize *IGF2BP2;* scPrediXcan and TWAS-bulk concurrently identify five silver-standard genes, underscoring scPrediXcan’s higher sensitivity in detecting T2D-related genes.

To investigate the cell-type specificity of the association with T2D risk, we analyzed the scPrediXcan results for all 11 islet cell types in the T2D dataset (Fig. 5e). We observed that while most TWAS hits were common across different cell types—48 (9.3%) genes are shared by all and 392 (76.1%) appeared in at least two cell types—123 (23.9%) genes are unique to one cell type.

To further examine the cell-type specificity of the 121 genes identified in only one cell type, we aggregated p-values across all remaining cell types using the ACAT method. After aggregation, two genes reached Bonferroni significance. While 118 genes were nominally significant (p < 0.05) in other cell types—these are referred to as cell-type–enriched genes. The remaining three genes (*CYB561*, *PISD*, and *RREB1*), referred to as cell-type–specific genes, did not reach nominal significance in any other cell type (supplementary fig. 3a). This suggests that associations involving cell-type–enriched genes may not be strictly cell-type–specific, but rather enriched in the focal cell type.

Our cell-type–enriched results also highlight several genes previously proposed as candidate drivers of T2D that were missed by TWAS-bulk analyses conducted on pancreas tissue, illustrating the advantages of performing TWAS at the cell-type level.

For example, the gene *CASR*, which mediates white adipose tissue dysfunction to promote the development of obesity-induced T2D^25^ reached significance only in gamma cells (p = 2.3 × 10^−6^); *MSRA,* which can cause oxidative stress when down-regulated leading to obesity-induced T2D^26^, is identified only in activated stellate cells (p = 1.6 × 10^−7^); and *LPL*, associated with a lower risk of T2D when up-regulated^27^, is found exclusively in quiescent stellate cells (p = 7.6 × 10^−8^). These findings highlight the potential of scPrediXcan to uncover nuanced, cell-type–enriched pathways involved in disease processes. The scPrediXcan T2D results for all cell types are provided in Supplementary Tables 4-14.

Finally, we benchmarked scPrediXcan, TWAS-pseudobulk, and TWAS-bulk against five other gene prioritization methods—Effector Index^28^, Polygenic Priority Score^29^, 3D chromatin^30^, CRISPR-screen^31^, and COLOC^32,33^—using the T2D ‘silver-standard’ genes to evaluate each method’s precision and recall (Methods). Among the eight gene prioritization methods, scPrediXcan demonstrates the second-highest recall at 0.439, surpassed only by the Effector Index method’s 0.561. Although scPrediXcan ranks 4th in precision (0.109), it is noteworthy that precision scores are generally low across all computational methods (0.046–0.175), except for the CRISPR-screen method (0.40). This pattern suggests that PrediXcan’s limited precision may stem from the silver-standard gene list not fully capturing the complex genetic landscape of T2D. The high recall rate of scPrediXcan highlights its robust ability to identify relevant genes, demonstrating its effectiveness despite the precision limitations of computational gene prioritization methods.

### scPrediXcan in systemic lupus erythematosus outperforms canonical TWAS methods by identifying promising candidate causal genes in more genomic loci

Next, we performed TWAS for systemic lupus erythematosus (SLE) using scPrediXcan and benchmarked its performance against the two canonical TWAS methods (pseudobulk and bulk, see Methods). scPrediXcan identifies a greater number of candidate causal genes and explains more GWAS loci for SLE than both the TWAS-pseudobulk and TWAS-bulk frameworks. Representative results for transitional B cells (scPrediXcan and TWAS-pseudobulk) and whole blood (TWAS-bulk) are shown in Figure 6a.

**Figure 6:**
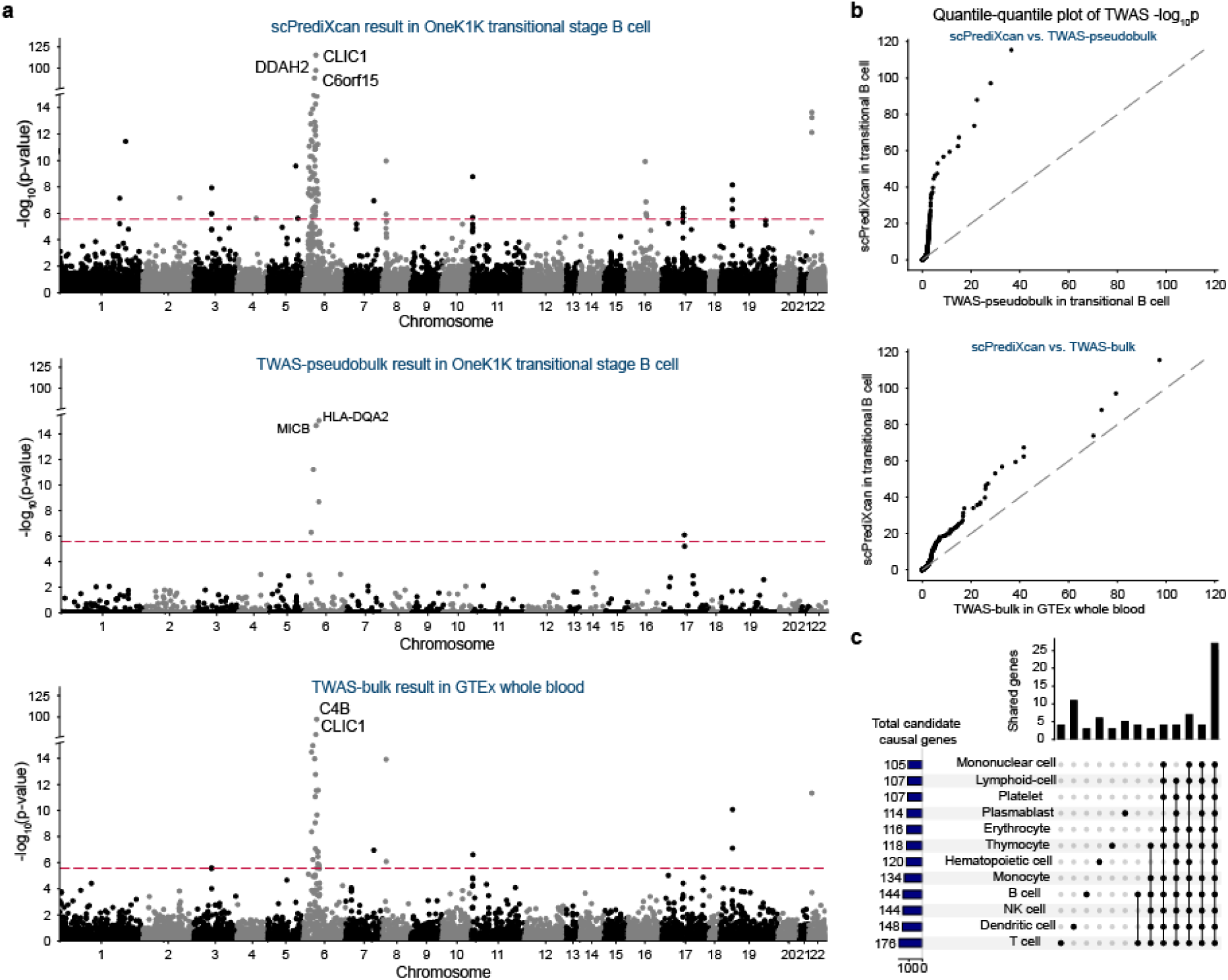
scPrediXcan in systemic lupus erythematosus outperforms canonical TWAS methods. **a**) Manhattan plots for different SLE TWAS frameworks. Top: scPrediXcan in transitional B cell from OneK1K dataset. Middle: TWAS-pseudobulk in transitional B cell from OneK1K dataset. Bottom: TWAS-bulk in whole blood from GTEx dataset. The red dashed lines are Bonferroni-corrected thresholds (p < 0.05/number of genes in the association study). **b**) QQ-plots of SLE TWAS p-values between frameworks. **c**) UpSet plot of scPrediXcan nominated candidate causal genes for SLE in different cell types.

To evaluate the effectiveness of our approach, we conducted a two-step comparison of the TWAS p-values. First, for genes predicted by all three models, we directly compared the p-values (supplementary fig. 6b). For a more fair comparison, we imputed genes that were not predicted by the canonical frameworks with uniformly distributed p-values. This comparison shows that scPrediXcan significantly outperforms the other methods (Fig. 5b).

Specifically, scPrediXcan identifies 129 candidate causal genes from 24 different LD blocks out of 1703 pre-defined LD blocks^23^, whereas TWAS-pseudobulk only identifies 11 candidate causal genes from 8 different LD blocks, and TWAS-bulk identifies 54 candidate causal genes from 14 different LD blocks. As expected, we noticed that both scPrediXcan and TWAS-bulk nominate many candidates at the HLA region on chromosome 6. scPrediXcan also pinpoints significant genes on other chromosomes that the other frameworks miss. For example, only scPrediXcan identifies two genes on chromosome 16 that have been implicated in SLE pathogenesis: *PYCARD* (also named *ASC*) in CD4+ alpha-beta T cell (p=7.8 × 10^−48^) and *ITGAM* in CD14+ monocytes (p=4.3 × 10^−41^)^34,35^.

To investigate the cell-type specificity of the association with SLE risk, we analyzed the scPrediXcan results of 12 immune cell types. To simplify interpretation, we aggregated the results of the various subtypes of T cells using the ACAT method. Similar to our findings for T2D, we found that while most TWAS hits are shared by different cell types, a few are cell-type–enriched, which we defined above as Bonferroni significant in one cell-type and nominally significant in others (Fig. 6c). Among 243 TWAS hits from the 12 immune cell types, 27 (11.1%) genes are shared in all cell types, 205 (84.3%) genes are shared in at least two cell types, and 38 (15.6%) genes yield significance in a single cell type. The full set of scPrediXcan SLE results are in Supplementary Tables 15–43.

To further investigate the cell-type specificity of the 38 genes identified in only one cell type, we aggregated p-values across all remaining cell types using the ACAT method. Our analysis revealed that 18 of these genes were cell-type–enriched, i.e., nominally significant (p < 0.05) in non-focal cell types. The remaining 20 genes were cell-type–specific, as the association did not reach nominal significance (p > 0.05) even after aggregation across all non-focal cell types (supplementary fig. 3b).

We identified several potential driver genes for SLE among the cell-type–specific and cell-type–enriched associations that were overlooked in the bulk TWAS analysis. One notable example is the complement factor B gene (*CFB*) identified by scPrediXcan as a cell-type–specific gene associated with SLE risk^36,37^ in T cells (p=2.8 × 10^−8^). Deficiencies in the classical complement pathway significantly contribute to SLE predisposition, as its disruption impairs the clearance of apoptotic cells and initiates an autoimmune response through the recognition of cellular debris by autoantibodies. This leads to a loss of tolerance by antigen-presenting cells and subsequent activation of T cells^37,38^. Importantly, *CFB* is part of the alternate complement pathway. Genes in these pathways are crucial for cellular clearance and immunity, and their upregulation has been reported in other autoimmune diseases like lupus nephritis^39^ suggesting potential therapeutic utility. Another example is *CXCR5*, identified in plasmablast cells (p = 1.4 × 10^−6^), has been found to be differentially expressed in SLE patients compared to healthy controls^40^. Notably, *CXCR5* was reported to be critically involved in the progression of lupus^41^. These examples showcase the ability of scPrediXcan to connect GWAS loci with genes known to have roles in SLE and other diseases, but that had been missed by canonical TWAS and other expression-based integrative approaches.

## Discussion

In this study, we presented scPrediXcan, a framework designed to perform transcriptome-wide association studies (TWAS) at the cell-type level. By leveraging transfer learning from a pre-trained deep-learning model and integrating single-cell expression data, we trained cell-type–specific gene expression predictors. We applied scPrediXcan to both T2D and SLE, benchmarking our results against canonical TWAS frameworks under various training conditions. Our findings indicate that scPrediXcan not only identifies a larger number of candidate causal genes and explains more GWAS loci, but also exhibits higher power in identifying candidate causal genes, especially when tested against a curated set of silver-standard T2D genes and genes with prior evidence of involvement in SLE. Moreover, scPrediXcan demonstrated enhanced sensitivity in nominating candidate causal T2D genes compared to other gene prioritization methods, such as COLOC, PoPS, and Effector index, among others.

Three key factors contribute to the enhanced performance of scPrediXcan: First, scPrediXcan utilizes a cross-genome deep learning model for gene expression prediction using reads aggregated across individuals, effectively reducing the impact of data sparsity and leveraging insights from a pre-trained sequence-to-epigenomics model. This strategy enables the prediction of a broader array of genes with high accuracy. Second, unlike the canonical TWAS framework, which typically operates at the tissue level, scPrediXcan focuses on the cell-type level. This granularity provides a more resolved context for identifying putative causal genes and captures those that might be overlooked at the tissue level. Third, scPrediXcan’s lower sample size requirement allows for the utilization of patient data that are often more disease-relevant, but less available, than data from healthy controls. This aspect of scPrediXcan is particularly advantageous as it can reveal candidate disease drivers that may remain hidden in non-disease contexts.

While scPrediXcan presents a robust framework for conducting TWAS at the cell-type level, there are certain limitations to consider. Some predictions correlate negatively with observed expression levels, a common challenge in deep learning models like ctPred that predict molecular phenotypes from reference genomes. Hence, our current analysis focuses on the p-values of correlations and associations and does not consider the direction of correlation. However, we are still limited in our ability to discern whether disease risk is associated with an up- or down-regulation of a nominated gene, a crucial piece of information for drug target selection and development. We are actively working to refine this aspect, with improvements anticipated as enhanced pre-trained models become available. Moreover, like all TWAS frameworks, scPrediXcan primarily focuses on cis-regulatory mechanisms and does not account for trans-effects or other regulatory mechanisms, and it is susceptible to linkage disequilibrium-induced errors.

In summary, scPrediXcan leverages single-cell RNAseq data and GWAS summary statistics to perform TWAS at the cell-type level, offering significant potential to identify putative causal genes in disease-relevant cell types. This capability advances our understanding of disease etiology and supports future experimental and clinical research.

To facilitate broad adoption, we make scPrediXcan and its integrated SNP-based linear predictors for 46 cell types publicly available at predictdb.org, providing a user-friendly tool for nominating context-specific causal genes. This resource can be used by end-users without expertise or infrastructure to perform deep learning analysis.

## Methods

### Data

#### Single-cell transcriptomics data and genotype data

The raw cell by gene matrices of OneK1K single-cell transcriptomics data12 and genotype data were shared by Dr. Joseph Powell. The data was pre-processed by Dr. Powell’s team described briefly as follows. The cell-droplets identified as doublets by both Demuxlet^42^ and Scrublet^43^ were removed. For each pool of cells captured, the distributions of total number of UMIs, number of genes, and percentage of mitochondrial gene expression were normalized using an Ordered Quantile Transformation. Then a generalized linear model with SCTransform^44^ method was used to account for batch effects and to get the gene UMI count matrix. Cells were classified into the major immune populations in a supervised manner using the gene expression data by a digital single cell transcriptional profiling panel^53^ as a reference. After the pre-processing steps done by Dr. Powell’s team, for further quality control, we selected cells with ‘nCount_RNA’ less than 10000 to avoid potential doublets and multilets, and the percentage of mitochondria RNA less than 10%. Then, we used the filtered data with its original cell type annotations reference for the downstream analysis.

The T2D single-cell transcriptomics data and genotype data were shared by Dr. Rohit Kulkarni. The data was pre-processed by Dr. Rohit Kulkarni’s team: the UMI count matrix was filtered by quality control requirements for cells to express at least 200 gene features and each gene feature to be present in at least three cells. Then doublets and triplets were removed using DoubletDecon^45^. Cells were manually annotated using a list of canonical markers from previous human islet single-cell RNA-seq studies. After the pre-processing steps done by Dr. Rohit Kulkarni’s team, the processed data with its cell type annotations was used for the downstream analysis.

The Tabula Sapiens single-cell transcriptomics data was downloaded online^14^. The data was pre-processed by the Tabula Sapiens Consortium. Briefly, cells that did not have at least 200 detected genes were removed, and then cells with fewer than 5000 counts and for droplet cells with fewer than 2500 UMIs were removed. DecontX^46^ was used to filter out reads from ambient RNA. The dataset was re-filtered by excluding the mitochondrial encoded genes when removing cells that did not contain the minimum number of genes and/or minimum of counts/UMIs to get the gene-count matrix. Cells were classified into different cell types using annotation methods include random forest (RF)^47^, support vector machine (SVM)^47^, scANVI^48^, onClass^49^, and k nearest neighbours (kNN) after batch-correction using single-cell harmonization methods (scVI^50^, BBKNN^51^, Scanorama^52^). After the preprocessing steps done by Tabula Sapiens team, we used the filtered gene-count matrix with cell annotations for the downstream analysis.

#### GWAS summary statistics

We used the multi-ancestry GWAS meta-analysis summary statistics^30^ from the DIAGRAM (DIAbetes Genetics Replication And Meta-analysis) consortium. We lifted over SNP coordinates to hg38 coordinates using UCSCs liftover tool to map variants between genome builds.

We downloaded the GWAS summary statistics of systemic lupus erythematosus^35^ from the GWAS Catalog.

### Computational methods

#### scPrediXcan framework

##### ctPred model training and prediction

###### Training data preprocessing

We used Enformer to predict the epigenomic features surrounding the TSS of each gene. We fed Enformer with a sequence length of 196,608 base pairs (bp) centered at the transcription-start-site (TSS) and generated an 896 by 5313 matrix representing the central 114,688 bp window. Within this matrix, each of the 896 bins contained one epigenomic feature for a 128-bp segment, encompassing a total of 5313 distinct features.

To reduce the computational burden for the training of ctPred, we averaged the central four bins (447-450th) as the local regions of gene TSS into a linear 1 by 5315 vector as the final epigenomic representation of the gene. We determined this method maintained the best prediction performance through empirical testing, The epigenomic representations of genes were used as the inputs for the ctPred model.

For each cell-type–specific gene-by-individual count matrix at pseudobulk level, we averaged the read counts of each gene across individuals, ranked the averaged counts for each gene across all individuals, and converted the ranks into percentiles ranging from 0 to 1. These percentile values were used as the target outputs for the ctPred model (Supplementary Fig. 8).

The minimum number of cells per cell type used for ctPred training was 125, with a corresponding minimum total read count of 561,372 reads per cell type. Although increasing the number of cells and total read counts generally improves ctPred prediction performance (Supplementary Fig. 7), the chosen thresholds for cell numbers and read counts were sufficient to maintain reasonable prediction accuracy.

###### Fully-connected neural network training

The model ctPred is a four-layer multi-layer perceptron (MLP) that predicts cell-type–specific gene expression levels given their epigenomic representations. The input is the reference epigenomic representation of a gene and the output is the rank-based gene expression percentile value for a specific cell type. Distinct models were trained for each cell type. The whole ctPred model consists of a linear layer that maps the 5313 dimensional epigenomic representation to a hidden layer of 64 dimensions, followed by ReLU, dropout, and other three identical hidden layers to map to the final predictions. We split the protein-coding genes into training (14429), validation (2812), and test (2426) sets by different chromosomes to avoid data leakage (Fig. 2a). The MSE (mean squared error) loss was used as the loss function of the model. To avoid overfitting, we used dropout layers (dropout rate = 0.05) and applied weight decay (L2 regularization parameter = 5 × 10^−4^). The model was saturated after 50–80 epochs. Finally, for model evaluation, the Pearson correlation between the observations and the predictions across genes in the test set was calculated.

###### Personalized gene expression prediction using ctPred

The personalized gene expression prediction process involves two inference steps: 1) Epigenomic Representation Inference: Personal genome sequences centered at each gene’s TSS are input into Enformer to generate personalized gene epigenomic representations. 2) Gene Expression Prediction: These personalized epigenomic representations are used by ctPred to obtain individual predictions of gene expression.

###### ctPred model linearization

We generated an in-silico cell-type–specific gene expression reference dataset by predicting gene expressions for 462 European individuals from the 1000 Genomes Project. Specifically, the DNA sequences centered at each gene’s transcription start site (TSS) for different individuals were input into Enformer to obtain personalized gene representations. These representations were then processed by ctPred to predict individual gene expressions, forming the in-silico reference dataset. Subsequently, we linearized ctPred into ℓ-ctPred by fitting the genotype data and in-silico gene expressions with an SNP-based elastic net model, following the standard PrediXcan pipeline^1^. For validation, we calculated the 10-fold cross-validated Spearman correlations between ctPred predictions and ℓ-ctPred predictions for all genes (Fig. 4).

###### Performing association test by S-PrediXcan

We used the GWAS summary statistics of T2D and SLE, as well as the ℓ-ctPred model for TWAS. We calculated the z-score introduced by the Summary-PrediXcan method^12^ to evaluate the associations between genes and the trait. The z-score of gene–trait association is calculated:

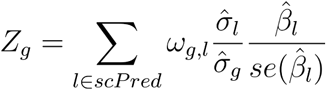

Where *ω_g,l_* is the effect of *SNP_l_* on the 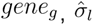 is the estimated variance of *SNP_l_*, and 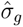 is the estimated variance of gene expressions of 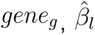 is the estimated effect size of *SNP_l_* on the trait and 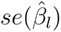 is the standard error of the effect size of 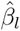. Th *ω_g,l_* is from the ℓ-ctPred model and the other statistics are calculated from GWAS summary statistics. Moreover, for the same gene–trait pair, a two-tailed p-value can be calculated from the z-score. All the scPrediXcan results for T2D and SLE in different cell types are attached in the supplementary tables.

### Canonical TWAS framework

The canonical TWAS frameworks serve as the benchmark against which we evaluate scPrediXcan’s performance.

#### TWAS-pseudobulk

##### Pseudobulk elastic net (PEN) training and prediction

The observed gene expression data at the pseudobulk level (i.e., a gene-by-individual count matrix for each cell type) was pre-processed using the same method as for ctPred. We averaged the counts for each gene across individuals, ranked the genes based on their average, and converted these ranks into percentiles ranging from 0 to 1. For each gene in a given cell type, we trained an SNP-based elastic net model by fitting the rank-based gene expression and the SNP-dosages at the cis-regions, following the standard PrediXcan pipeline.

To compare the PEN model against ctPred for personalized prediction in the T2D dataset, we divided the 29 individuals into a training set of 20 and a test set of 9. For downstream TWAS analysis, we used all 29 individuals for model training. In the OneK1K dataset, for model comparison against ctPred for personalized prediction, we randomly selected 800 individuals as the training set and 100 individuals as the test set. For downstream TWAS analysis, we used all individuals for model training.

##### Performing association test by S-PrediXcan

We followed the same workflow as the scPrediXcan association test using the S-PrediXcan method introduced earlier.

#### TWAS-bulk

##### SNP-based elastic net

For T2D, we directly downloaded the SNP-based elastic net models trained by GTEx pancreas bulk gene expression and genotype data (n = 305). For SLE, we directly downloaded the SNP-based elastic net models trained by GTEx whole blood bulk gene expression and genotype data (n = 670).

##### Performing association test by S-PrediXcan

We followed the same workflow as the scPrediXcan association test using the S-PrediXcan method introduced earlier.

### T2D-associated gene prioritization methods comparison

For scPrediXcan and TWAS-pseudobulk frameworks, we applied ACAT p-value combination method^21^ to integrate the TWAS results of all the 11 cell types from islet in T2D dataset and used Bonferroni-corrected p values (p < 2.5 × 10^−6^ for scPrediXcan, p < 1.4 × 10^−5^ for TWAS-pseudobulk) as the threshold to obtain the final prioritized gene lists. For TWAS-bulk, we directly used Bonferroni-corrected p-values (p < 8.5 × 10^−6^) as the threshold to obtain the final prioritized gene list from GTEx pancreas tissue. For other methods, the prioritized gene lists were downloaded from the CMDKP database. The silver-standard T2D-associated gene list was used to calculate the precision and recall of all the methods. The precision was defined as the number of nominated silver-standard genes divided by the total number of nominated genes, and the recall was defined as the number of nominated silver-standard genes divided by the total number of silver-standard genes.

## Code availability

The code for scPrediXcan is available at https://github.com/hakyimlab/scPrediXcan. Prediction models are available at https://predictdb.org.

## Acknowledgements

Figure 1 was created using Biorender.com.

## Computing resources

This research used resources of the Argonne Leadership Computing Facility, which is a DOE Office of Science User Facility supported under Contract DE-AC02-06CH11357. This work was completed in part with resources provided by the University of Chicago’s Research Computing Center and Beagle3. We also acknowledge resources from the Center for Research Informatics, funded by the Biological Sciences Division at the University of Chicago, with additional funding provided by the Institute for Translational Medicine, CTSA grant number 2U54TR002389-06 from the National Institutes of Health.

## Funding Sources

YZ, MC, and HKI were partially funded by R01 GM126553 and R01 HG011883; YZ, SSM, TA, and HKI were funded in part by R01AA029688; HKI was in part funded by P30DK020595.

## Author information

**Committee of Genetic, Genomics, and Systems Biology, University of Chicago, Chicago, Illinois, United States of America**

Yichao Zhou, Temidayo Adeluwa, Saideep Gona

**Department of Medicine, Section of Genetic Medicine, University of Chicago, Chicago, Illinois, United States of America**

Hae Kyung Im, Mengjie Chen, Lisha Zhu, Sarah Sumner, Sofia Salazar-Magaña

**Department of Human Genetics, University of Chicago, Chicago, Illinois, United States of America**

Festus Nyasimi

**Data Science and Learning Division, Argonne National Laboratory, Chicago, Illinois, United States of America**

Ravi Madduri

**Department of Medicine, Harvard Medical School, Boston, Massachusetts, United States of America**

Rohit Kulkarni, Hyunki Kim

**Department of Pharmacy and Pharmaceutical Sciences, National University of Singapore, Singapore**

Boxiang Liu

**UNSW Cellular Genomics Futures Institute, University of New South Wales, Sydney, Australia**

Joseph Powell

## Declaration of Interests

Authors declare no conflict of interests.

## Author contributions

Conceptualization, Y. Z, M. C., and H. K. I.; methodology, Y. Z., T. A., S. G., and F. N.; software: Y. Z., T. A., S. G., and F. N.; formal analysis, Y. Z, and H.K.I.; data curation: Y. Z. and F. N.; Visualization: Y. Z. and S.S-M.; resources, M. C., R. M., R. K., and H. K.; writing – original draft, Y. Z., and S.S.; writing – review & editing, Y. Z., S.S., S.S-M., B. L., M. C., and H. K. I.; funding acquisition, M. C. and H. K. I.

## Supplementary information

### Supplementary Figures

**Supplementary fig. 1:**
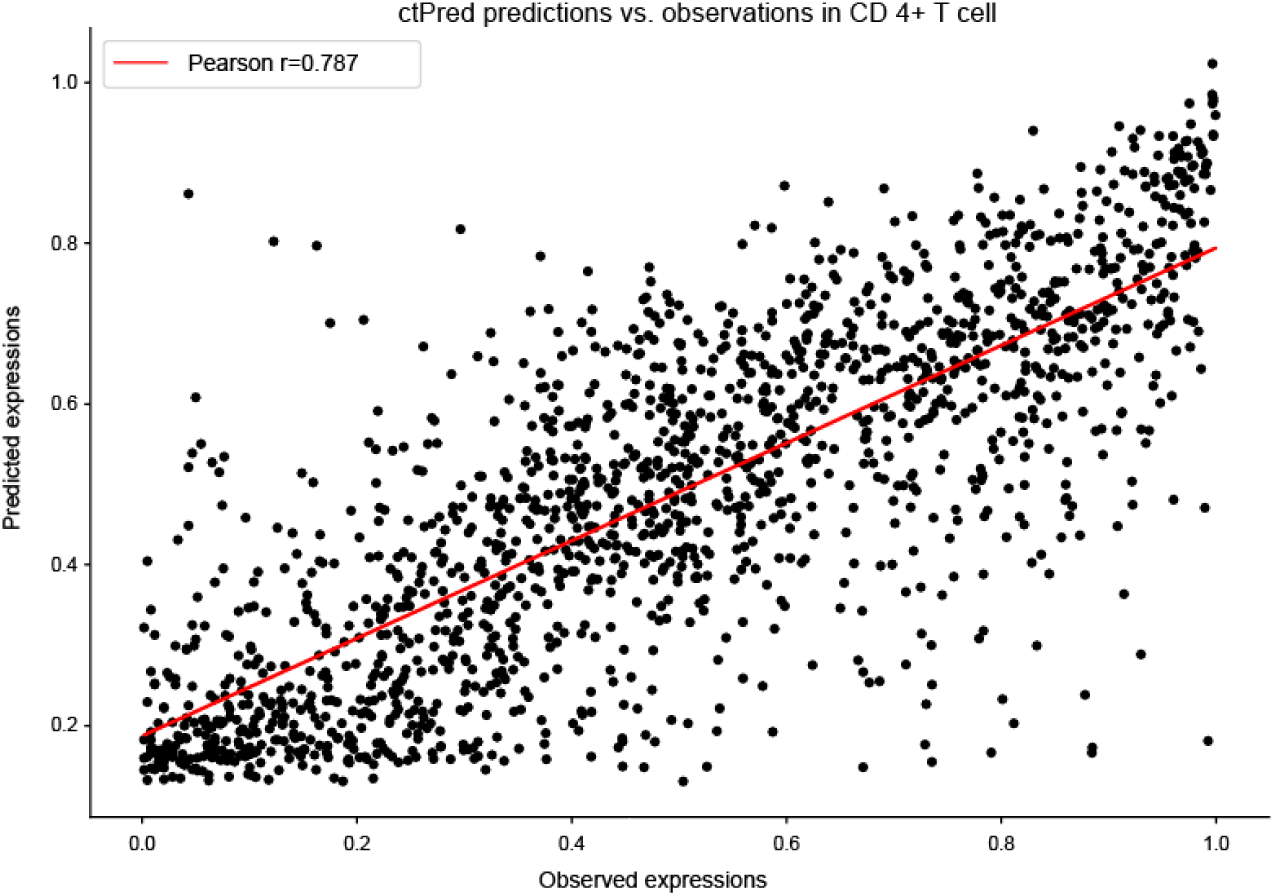
ctPred predicts cell type-specific gene expression in CD 4+ T cell. Scatter plot of ctPred predictions and observations for gene expressions in CD4+ T cell dataset.

**Supplementary fig. 2:**
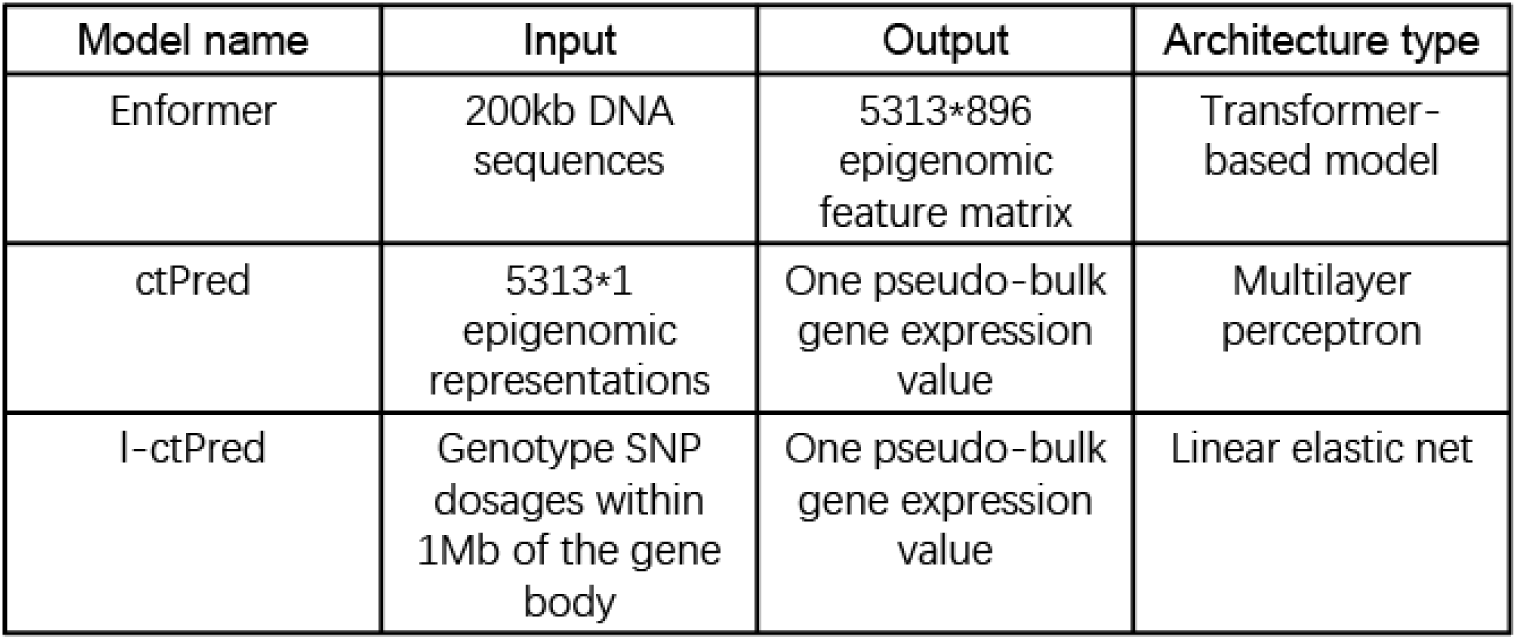
Brief description of the models in the scPrediXcan framework.

**Supplementary fig. 3:**
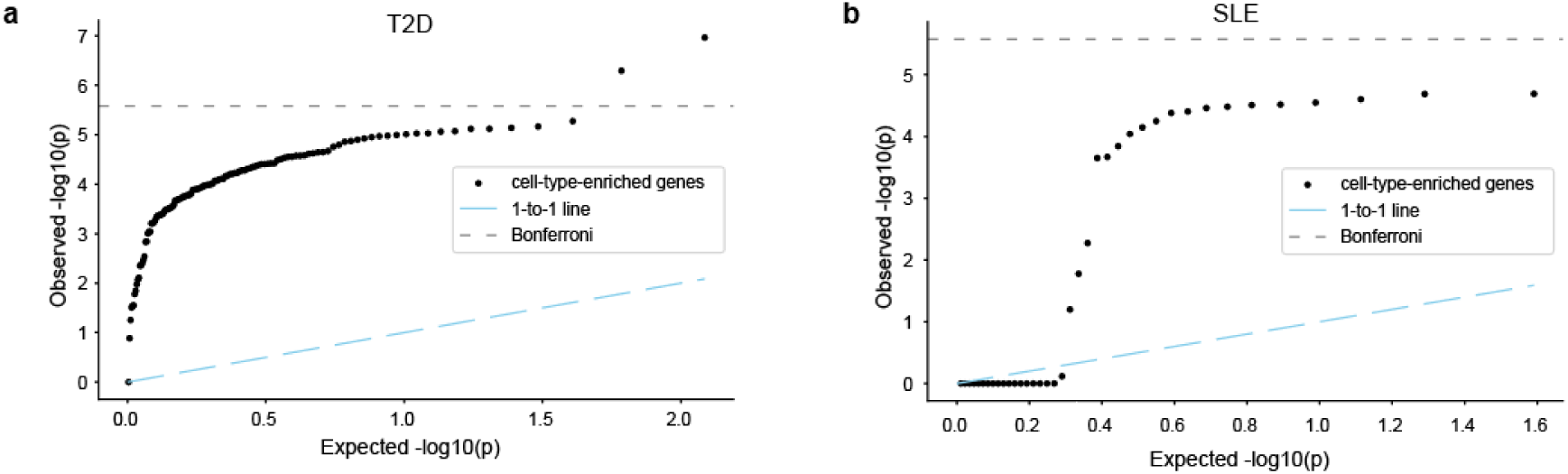
Quantile-quantile plot of ACAT-adjusted TWAS −log_10_ (p-value) against uniformly distributed p-value for T2D and SLE. **a)** Quantile-quantile plot of ACAT-aggregated TWAS −log10 (p-value) in all non-significant cell types for genes passing the Bonferroni-corrected threshold in only one islet cell type from T2D dataset for T2D trait. **b)** Quantile-quantile plot of ACAT-aggregated TWAS −log10 (p-value) in all non-significant cell types for genes passing the Bonferroni-corrected threshold in only one immune cell type from OneK1K dataset for SLE trait.

**Supplementary fig. 4:**
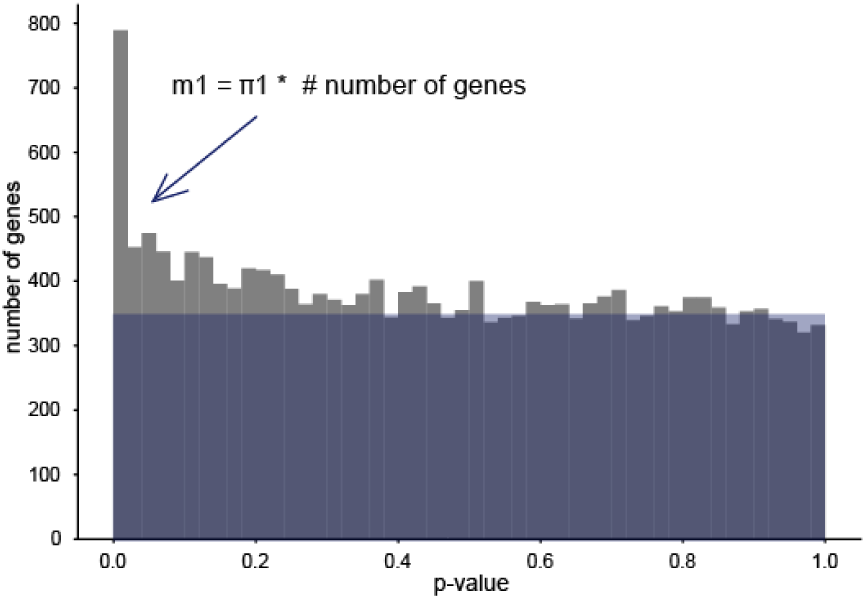
Calculating the number of true positive genes with p-values deviating from a uniform distribution. The m1 calculation is based on the histogram of p-values of Pearson correlations between predicted gene expressions and observed gene expressions. The blue rectangular region shows the genes with p-values following a uniform distribution, and the proportion of those genes is denoted as pi0. The pi1 is 1-pi0, and the number of true positive genes is calculated by pi1 * total number of genes.

**Supplementary fig. 5:**
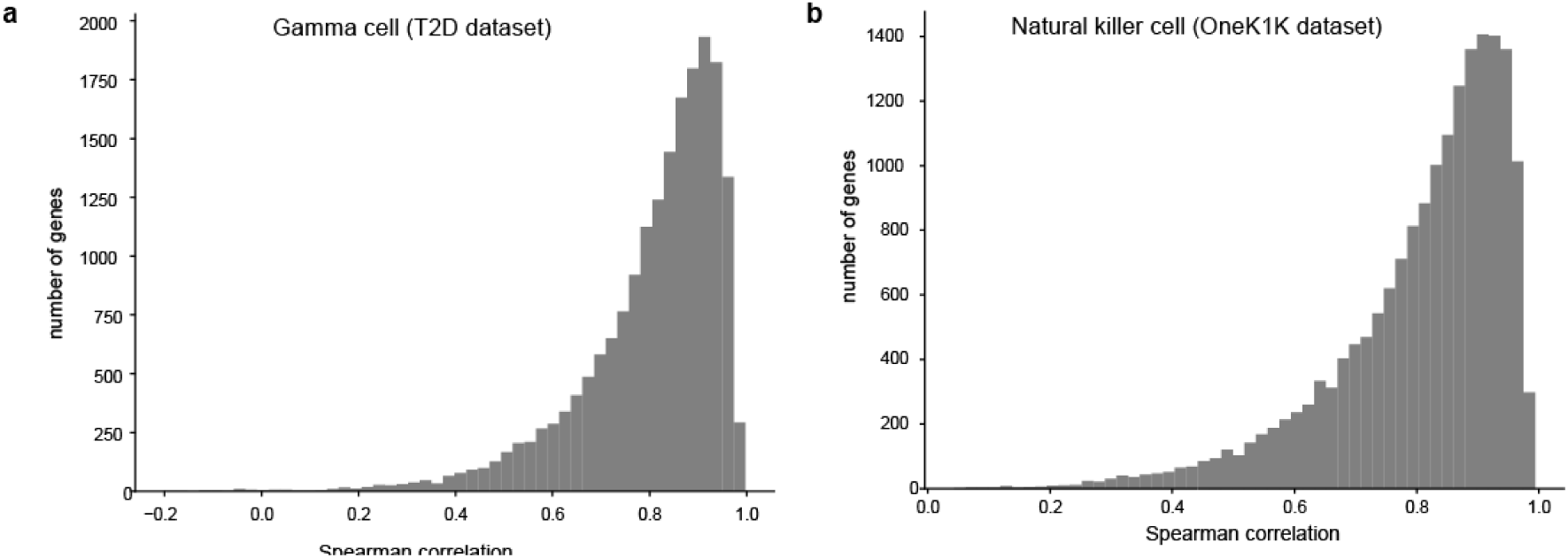
Distribution of Spearman correlations between ctPred predictions and l-ctPred predictions of all genes in representing cell types. **a)** Histogram of Spearman correlations between ctPred-predicted gene expressions and l-ctPred-predicted gene expressions in gamma cell as the representing cell type from T2D dataset. Other cell types have similar correlation distributions. **b)** Histogram of Spearman correlations between ctPred-predicted gene expressions and l-ctPred-predicted gene expressions in natural killer cells as the representing cell type from OneK1K dataset. Other cell types have similar correlation distributions.

**Supplementary fig. 6:**
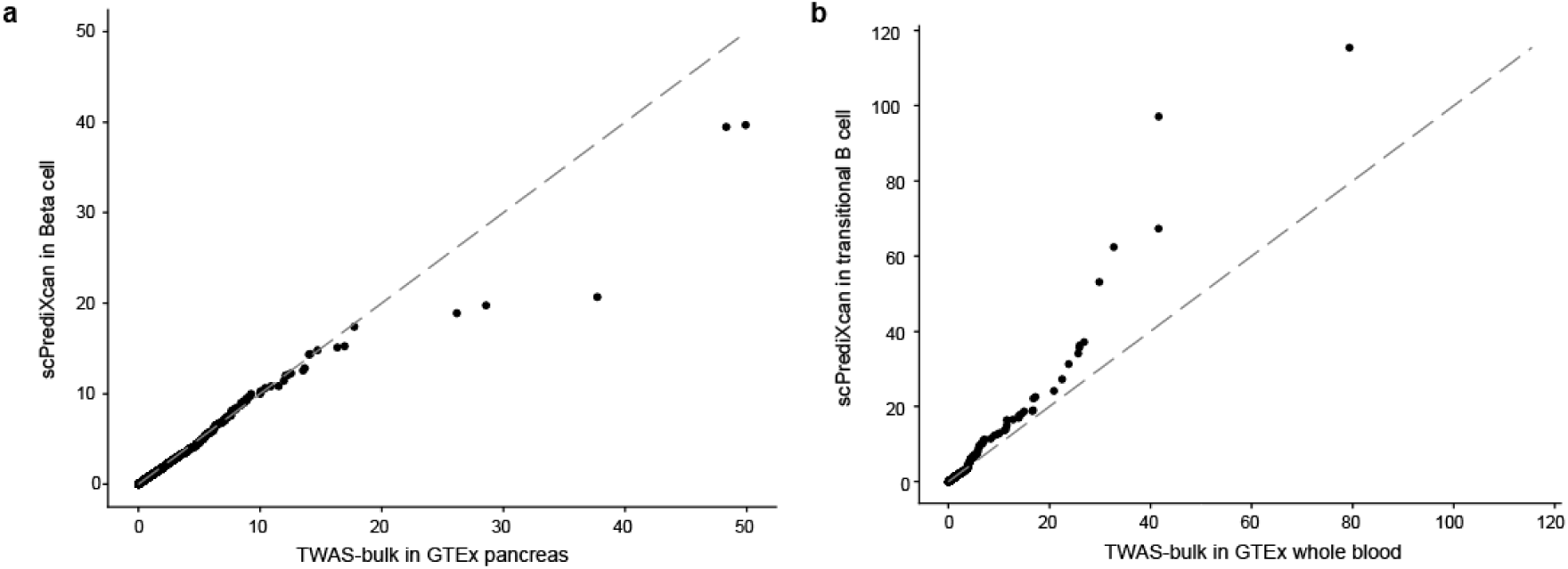
Quantile-quantile plot of TWAS −log_10_ (p-value) for only the overlapped genes between models. **a)** Quantile-quantile plot of T2D TWAS −log10(p) of overlapped genes in scPrediXcan in Beta cell and TWAS-bulk in GTEx pancreas. This set of genes will likely favor the TWAS-bulk method since only models that performed well enough in this approach end up included here. A more fair comparison is shown in Figure 5b where union of genes tested by scPrediXCan and TWAS-bulk in GTEx pancreas are shown, imputing the p-values of genes missed by TWAS-bulk with uniformly distributed p-values. **b)** Quantile-quantile plot of SLE TWAS −log10(p) of overlapped genes in scPrediXcan in transitional B cell and TWAS-bulk in GTEx blood.

**Supplementary fig. 7:**
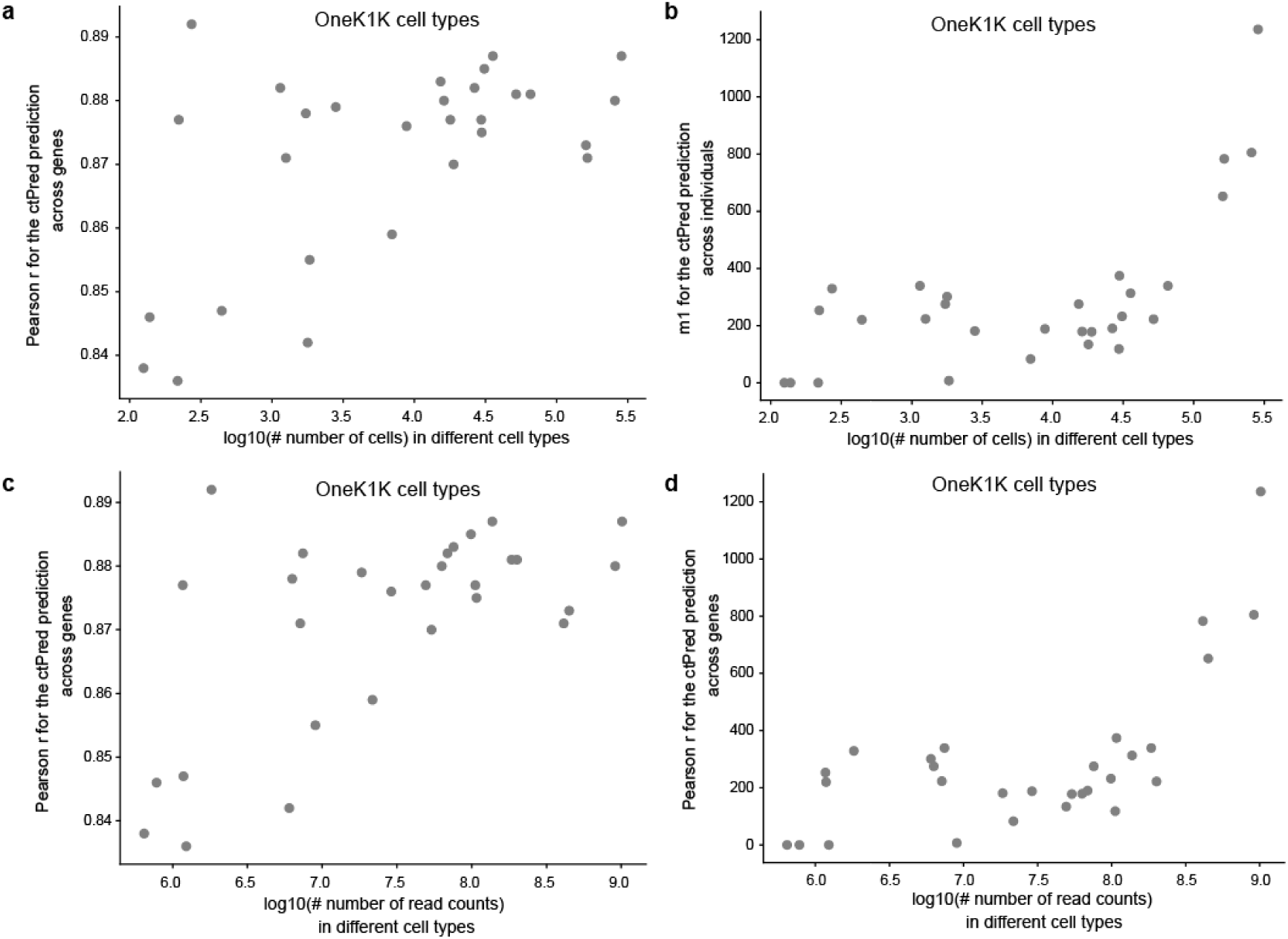
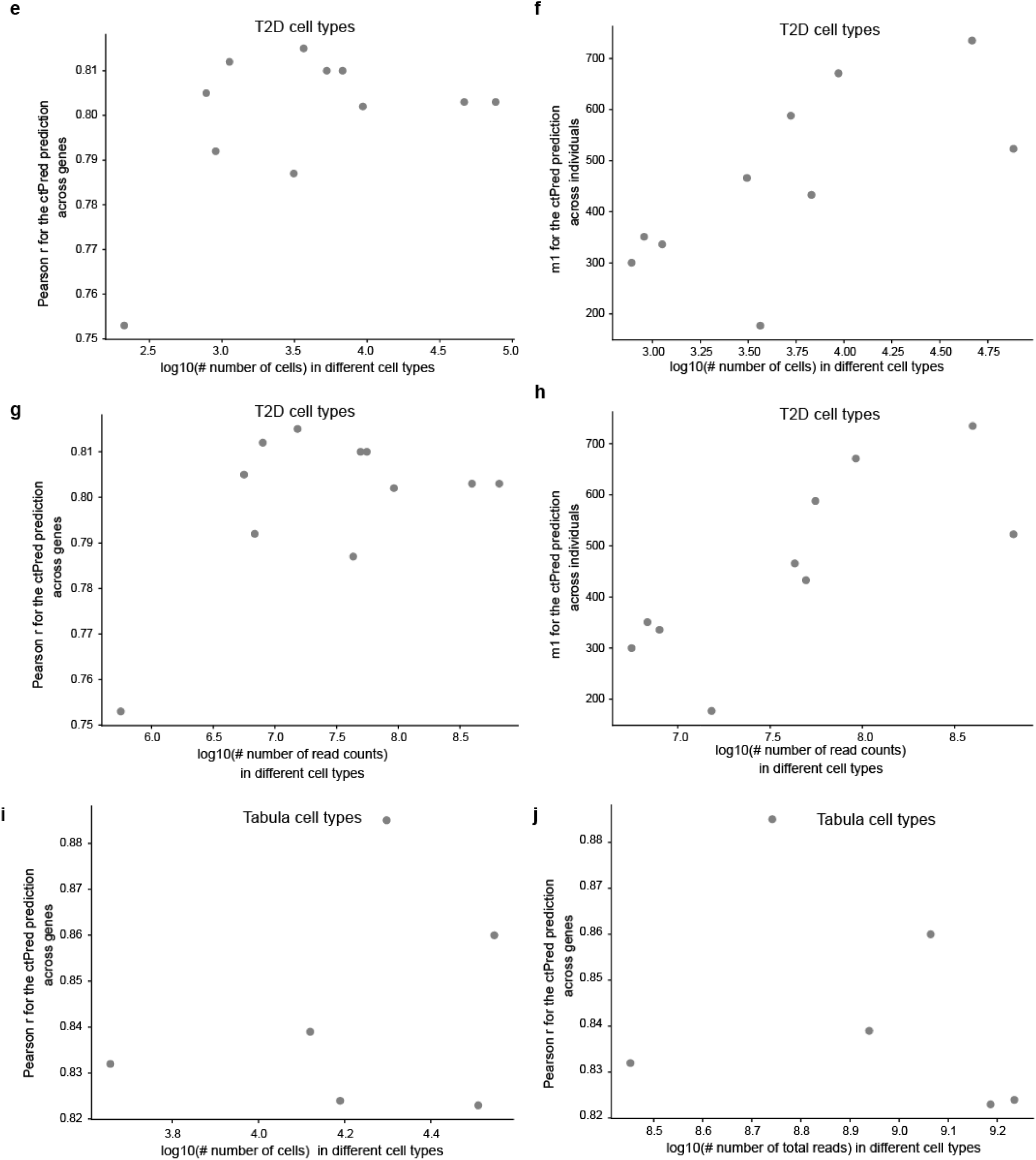
ctPred prediction performance metrics vs. cell numbers and read counts of different cell types across datasets. **a)-d) Prediction performance vs number of cells and read counts in OneK1K dataset.** Scatter plot of ctPred prediction metrics (Pearson r for prediction across genes or m1 value for prediction across individuals) and cell numbers or total scRNAseq read counts in different cell types from OneK1K dataset. See supplementary table tab. 44. **e)-h) Prediction performance vs number of cells and read counts in T2D dataset.** Scatter plot of ctPred prediction metrics (Pearson r for prediction across genes or m1 value for prediction across individuals) and cell numbers or total scRNAseq read counts in different cell types from T2D dataset. See supplementary table tab. 44. **i)-j) Prediction performance vs number of cells and read counts in Tabula Sapiens dataset.** Scatter plot of ctPred prediction metrics (Pearson r for prediction across genes for prediction across individuals) and cell numbers or total scRNAseq read counts in different cell types from Tabula Sapiens dataset. See supplementary table tab. 44.

**Supplementary fig. 8:**
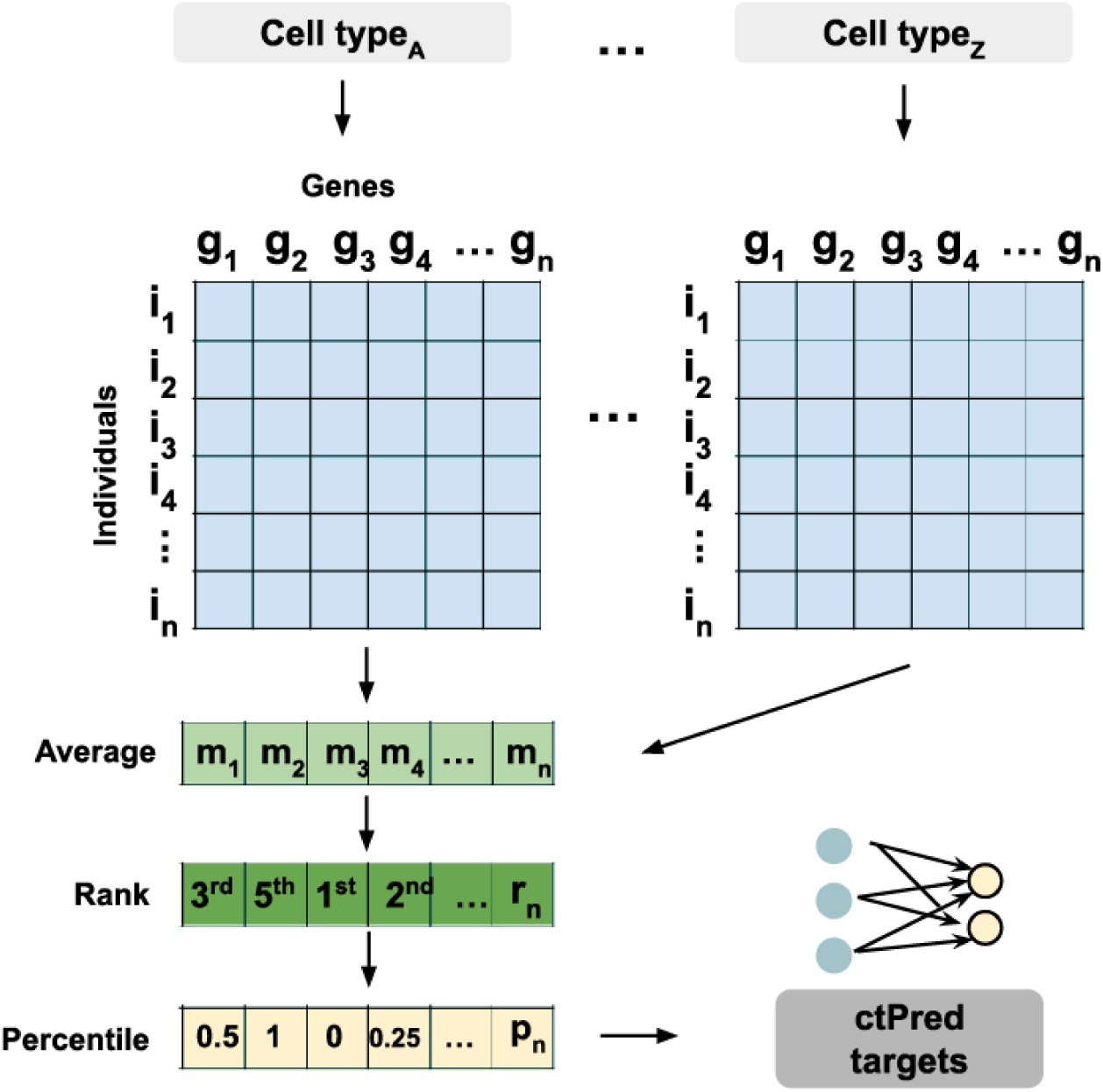
scRNA-seq pseudobulk data processing for ctPred training. The scRNA-seq processing into the target for ctPred model training.

### Supplementary tables

Supplementary tables 1: General information (tables 1-3, tables 44-45). Table 1: cell types in T2D, OneK1K and Tabula Sapiens datasets. Table 2: number of genes trained and converged of PEN in the canonical TWAS framework. Table 3: ctPred prediction performance across genes in different cell types of three datasets. Table 44: ctPred prediction performance against the number of cells or total read counts per cell type for training in different cell types from T2D, OneK1K and Tabula Sapiens datasets. Table 45: Gene names of T2D silver-standard genes.

Supplementary tables 2: T2D TWAS results (tables 4-14). Table 4-14: T2D TWAS association z-score, effect sizes and p-values of tested genes in different cell types from T2D dataset.

Supplementary tables 3: SLE TWAS results (tables 15-43). Table 15-43: SLE TWAS association z-score, effect sizes and p-values of tested genes in different cell types from OneK1K dataset.

## Notes

### Competing Interest Statement

The authors have declared no competing interest.

### Summary of Updates

A typo of the author's name has been corrected; Figure 4 revised with typo of legends corrected; Figure 5 revised with x coordinates corrected; Figure 6 revised with x coordinates corrected; Acknowledgement revised.

