## Supplementary material for "scPrediXcan integrates advances in deep learning and single-cell data into a powerful cell-type–specific transcriptome-wide association study framework": Sup_fig2.pdf

**Supplementary fig. 2: Brief description of the models in the scPrediXcan framework.**

| Model name | Input | Output | Architecture type |
|---|---|---|---|
| Enformer | 200kb DNA sequences | 5313*896 epigenomic feature matrix | Transformer-based model |
| ctPred | 5313*1 epigenomic representations | One pseudo-bulk gene expression value | Multilayer perceptron |
| l-ctPred | Genotype SNP dosages within 1Mb of the gene body | One pseudo-bulk gene expression value | Linear elastic net |
