## Supplementary material for "scPrediXcan integrates advances in deep learning and single-cell data into a powerful cell-type–specific transcriptome-wide association study framework": Sup_fig3.pdf

**Supplementary fig. 3: Quantile-quantile plot of ACAT-adjusted TWAS -$log_{10}$ (p-value) against uniformly distributed p-value for T2D and SLE.**

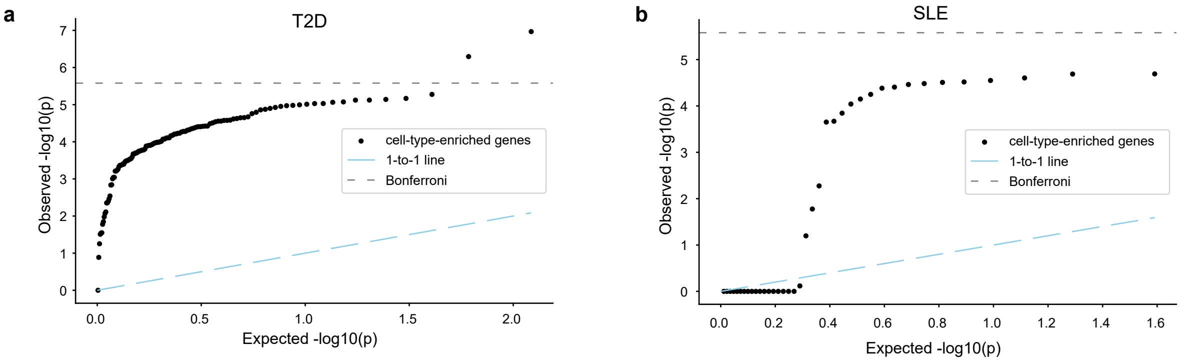
