## Supplementary material for "scPrediXcan integrates advances in deep learning and single-cell data into a powerful cell-type–specific transcriptome-wide association study framework": Sup_fig4.pdf

### Supplementary fig. 4: Calculating the number of true positive genes with p-values deviating from a uniform distribution.

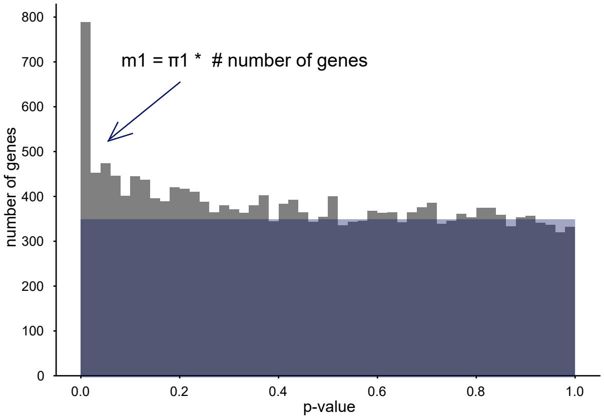
