## Supplementary material for "scPrediXcan integrates advances in deep learning and single-cell data into a powerful cell-type–specific transcriptome-wide association study framework": Sup_fig5.pdf

### Supplementary fig. 5: Distribution of Spearman correlations between ctPred predictions and l-ctPred predictions of all genes in representing cell types

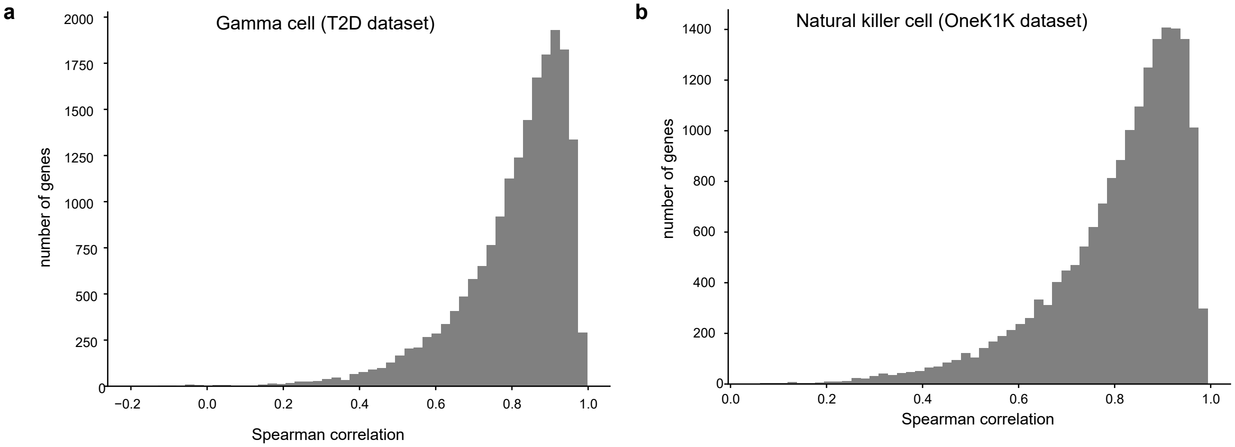
