## Supplementary material for "scPrediXcan integrates advances in deep learning and single-cell data into a powerful cell-type–specific transcriptome-wide association study framework": Sup_fig6.pdf

### Supplementary fig. 6: Quantile-quantile plot of TWAS -$\log_{10}$ (p-value) for only the overlapped genes between models

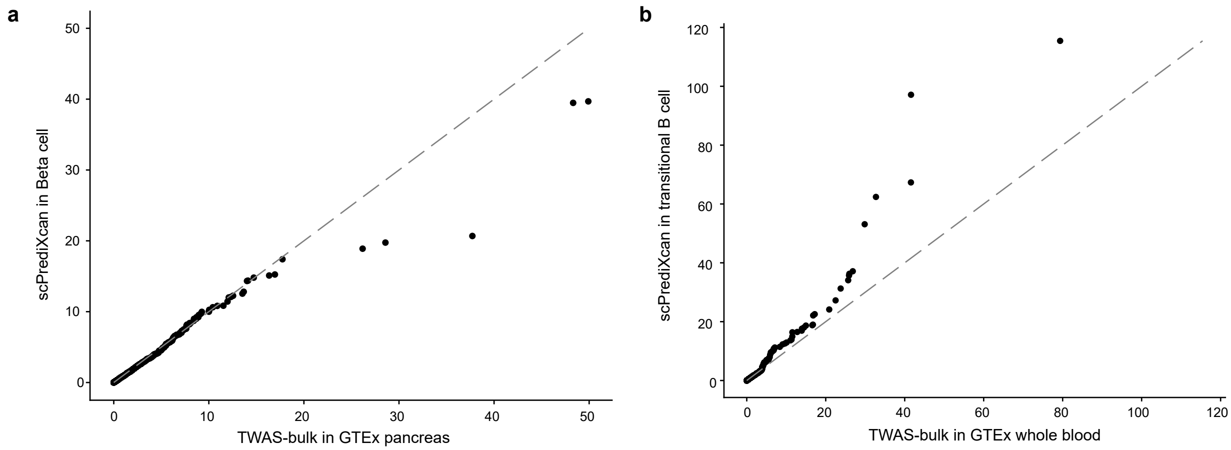
