## Supplementary material for "scPrediXcan integrates advances in deep learning and single-cell data into a powerful cell-type–specific transcriptome-wide association study framework": Sup_fig7.pdf

**Supplementary fig. 7: ctPred prediction performance metrics vs. cell numbers and read counts of different cell types across datasets.**

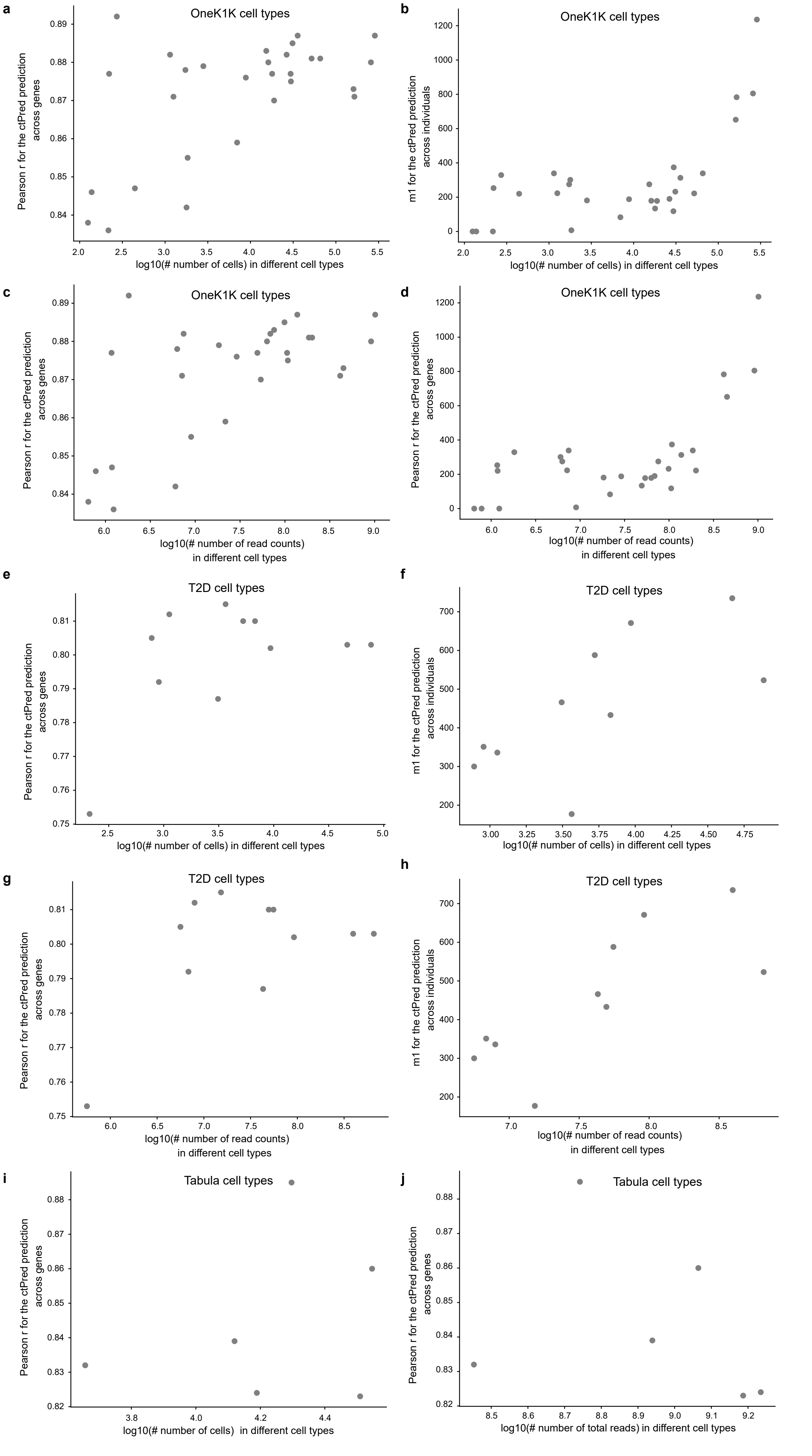
