## Supplementary figures and images for "scPrediXcan integrates advances in deep learning and single-cell data into a powerful cell-type–specific transcriptome-wide association study framework"

### Sup_fig1.pdf

#### Supplementary fig. 1: ctPred predicts cell type-specific gene expressions in CD 4+ T cell

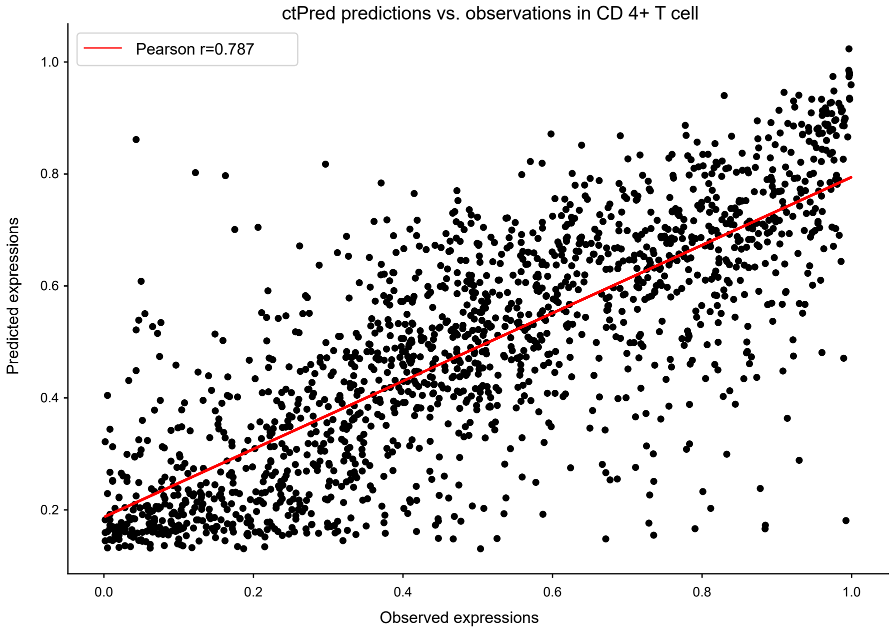

### Sup_fig8.pdf

**Supplementary fig. 8: scRNA-seq pseudobulk data processing for ctPred training.**

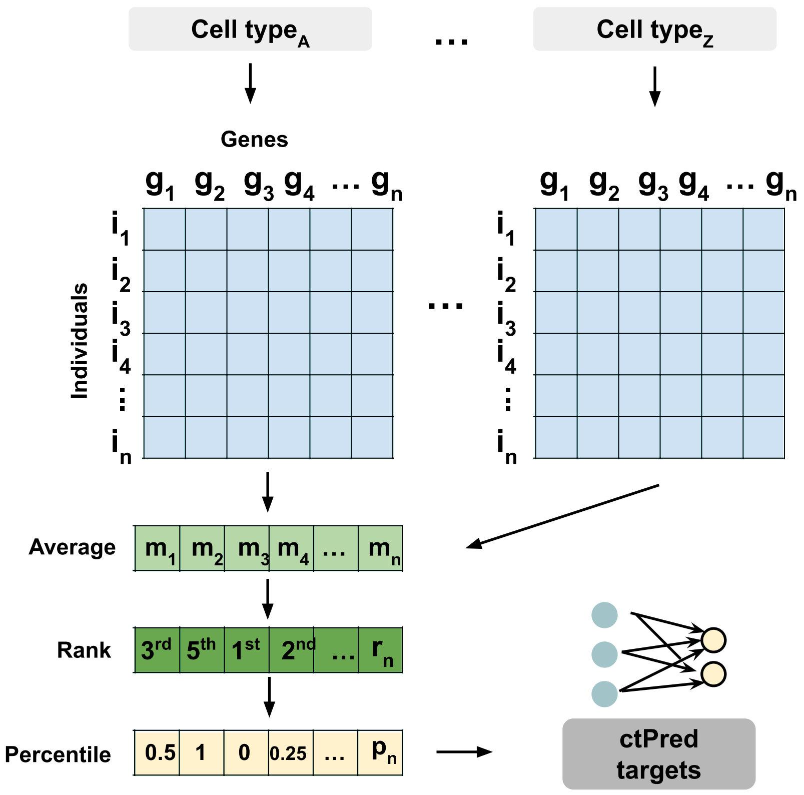
